# PML and PML-like exonucleases restrict retrotransposons in jawed vertebrates

**DOI:** 10.1101/2022.07.29.501749

**Authors:** Sabateeshan Mathavarajah, Kathleen L. Vergunst, Shelby K. Williams, Raymond He, Maria Maliougina, Elias B. Habib, Mika Park, Jayme Salsman, Stéphane Roy, Ingo Braasch, Andrew J. Roger, David N. Langelaan, Graham Dellaire

## Abstract

We have uncovered a novel role for the promyelocytic leukemia (*PML*) gene and novel PML-like DEDDh exonucleases in the maintenance of genome stability through the restriction of LINE-1 (L1) retrotransposition in jawed vertebrates. Although the PML tumour suppressor protein in mammals is SUMOylated and forms nuclear bodies, we found that the spotted gar PML ortholog and related proteins in fish are not SUMOylated and function as cytoplasmic DEDDh exonucleases. In contrast, more closely related avian and turtle PML proteins are predicted to be SUMOylated and localized both to the cytoplasm and to nuclear bodies. We also identified PML-like exon 9 (*Plex9*) genes in teleost fishes that encode exonucleases sharing homology to gar PML. In an example of convergent evolution and akin to TREX1, gar PML and zebrafish Plex9 proteins suppressed L1 retrotransposition and could complement *TREX1* knockout in mammalian cells. We also characterized the first non-mammalian TREX1 homologs in axolotl. Following export to the cytoplasm, the human PML-I isoform also restricted L1 through its conserved C-terminus and suppressed CGAS activation. Thus, PML first emerged as a cytoplasmic suppressor of retroelements, and this function is retained in amniotes despite its role in the assembly of nuclear bodies and the acquisition of SUMO-modification.

## INTRODUCTION

The *promyelocytic leukemia* (*PML*) gene was initially discovered as the oncogenic fusion partner of RARα in a translocation event that causes acute promyelocytic leukemia (1,2). In mammals, the PML protein forms a subnuclear organelle known as the PML nuclear body (NB) (1-3) that plays a prominent role in antiviral immunity and is a hub for the association and post-translational modification of over 150 proteins involved in transcriptional regulation, replication, and repair of DNA (3-5). PML is believed to contribute to tumour suppression through several mechanisms including the modulation of innate immune pathways as well as the regulation of the PTEN-AKT-mTOR signaling axis (3,6-9). Maintenance of genomic stability is also implicated in tumour suppression (10,11), and PML loss is associated with increased DNA damage and genome instability (5,8,12). Activation of the DNA damage response (DDR) triggers PML NB fission and the association of DNA repair factors with bodies, and loss or over-expression of PML inhibits DNA repair by homologous recombination (5,13,14). PML NBs also help maintain telomeric chromatin integrity in embryonic stem cells (15). However, a direct role for the PML protein itself in genome stability has remained elusive.

Endogenous retroelements such as the long-interspersed element-1 (LINE1/L1), are major contributors to genome instability and evolution (16,17).L1 elements are active autonomous retrotransposons, and almost one fifth of the human genome (17%) is comprised of L1 retroelement DNA (18). On average, human genomes contain 80-100 copies of retrotransposition-competent L1s, and these elements are responsible for the majority of our retrotransposition events (19-21). While L1s contribute a source of genetic variation, this is juxtaposed to how they corrupt genome integrity (17,22-24). For this reason, numerous host factors have been described as potent enhancers and suppressors of L1 activity (25,26). In addition to gene disruption by their insertion, L1 activation manifests in human disease through overstimulation of the DNA-sensing innate immune signalling pathways such as the CGAS-STING axis (27-29). Recent evidence has linked L1s as a driving force in autoimmune disease (29-34). For example, multiple genes associated with Aicardi-Goutières syndrome, an autoimmune disease sharing features with systemic lupus erythematosus, have been shown to encode regulators of L1 activity including *SAMHD1, ADAR1*, and *RNASEH2* (32,33,35-37). Aicardi-Goutières syndrome is also caused by loss of DEDDh family *Three-prime repair exonuclease 1* (*TREX1*), a potent suppressor of L1 activity and an essential brake in the type I interferon responses that when impaired contributes to systemic autoimmunity (38). Although TREX1 is the predominant DNA exonuclease in mammals, non-mammalian TREX1 homologs have not been characterized in other vertebrates, even in species whose genomes encode both L1 and the related L2 retrotransposons such as the teleost fishes (39).

Teleost bony fish species genomes including zebrafish lack a *PML* ortholog, presumably as a consequence of the tetraploidization of their genomes sometime between 300-450 million years ago (40,41). In mammals, the functional diversity of the human PML protein is derived from its 7 isoforms (PML-I to VII) that diverge at the C-terminus and are linked to specific cellular functions and protein interactions (14). However, most species of vertebrates including rodents, encode only a few PML isoforms, of which the most highly expressed resemble orthologs of the longest human isoform, PML-I. The human PML-I isoform is distinguished from other isoforms by encoding both a nuclear exclusion signal (NES) and an uncharacterized C-terminal domain encoded by exon 9 that shares homology to the DEDDh family of exonucleases and in particular, TREX1 (42,43). Here, we use a molecular evolution approach to better understand the conserved function of PML-I in vertebrate genome stability, uncovering the convergent evolution of novel PML-like DEDDh exonucleases in different fish lineages and amniotes, as well as a primordial role for PML in the suppression of L1 retrotransposition, a known driver of autoimmune disease and genome instability in cancer (44,45).

## MATERIAL AND METHODS

### Phylogenetic analyses

Two rounds of searches on the protein and transcriptome sequences were carried out to detect homologs to the human PML sequence in sequenced eukaryote species. A phylogenetic dataset of PML proteins was constructed by collecting 72 protein sequences from NCBI via a PSI-BLASTp or tBLASTn search followed by reverse BLASTP. For searches, in the first round we targeted both the full length PML (sequence for isoform PML-I) and the C-terminus of PML (an amino acid stretch from positions 600-882; the CDE or C-terminal DEDDh exonuclease domain) in the second round. The logic behind this stems from the RBCC motif being highly conserved and associated with TRIM19 proteins. The C-terminus is unique to PML. The *E-value* threshold (-e) was set as 0.001, the *E-value* threshold for inclusion in the multipass model (-h) as 0.002 (default value), and the maximum running iterations (-j) as 5. For transcriptome sequences, tBLASTn was used and the *E-value* threshold (-e) was set as 0.001. For species with fully sequenced genomes available, PML orthology was confirmed by synteny to the human *PML* loci.

A separate dataset of the Plex and TREX exonucleases was constructed through a similar approach but searching for both the C-terminus of PML, zebrafish Plex9.1 and Plex9.2 and TREX1. resulting in a dataset of 104 proteins. Previously identified homolog PML-CDE sequences that had homologous sequence and synteny were added to the Plex and TREX dataset. Both datasets were aligned using mafft einsi with default settings (46). The ExoIII dataset was trimmed so that only the ExoIII, and ExoIII-like, domains remained, while the C-terminal exonuclease domain was removed from the PML alignment. Both datasets were further trimmed using trimal v1.4.rev15 with the -gappyout setting (47), resulting in a PML data set containing 545 sites and a ExoIII dataset containing 242 sites. These datasets were used to estimate maximum-likelihood phylogenies using IQTree v1.5.5 (48) using the LG+C60+gamma model of evolution.

### PML RBCC domain phylogeny

The resulting PML phylogeny (Figure 1), created using the RBCC domains of the protein, shows the expected relatedness between major groups of metazoans, with moderate support. This analysis identifies the ancestral PML originating from within Chondrichthyes (the sharks & rays). This phylogeny highlights the divergence of the PML ortholog found in *Lepisosteus oculatus*, an Actinopterygii fish identified possessing a full-length PML sequence, which, as expected, is sister to Sarcopterygii. To assess domain architectures, InterPro 88.0 (https://www.ebi.ac.uk/interpro/) was utilized. Orthologs used in the analysis include ones from the following species (corresponds to silhouettes in Figure 1): *Amblyraja radiata, Scyliorhinus canicular, Lepisosteus oculatus, Tachyglossus aculeatus, Sarcophilus harrisii, Mus musculus, Pteropus vampyrus, Capra hircus, Ursus maritimus, Homo sapiens, Alligator sinensis, Pogona vitticeps, Chelonia mydas* and *Coturnix japonica*.

**Figure 1.**
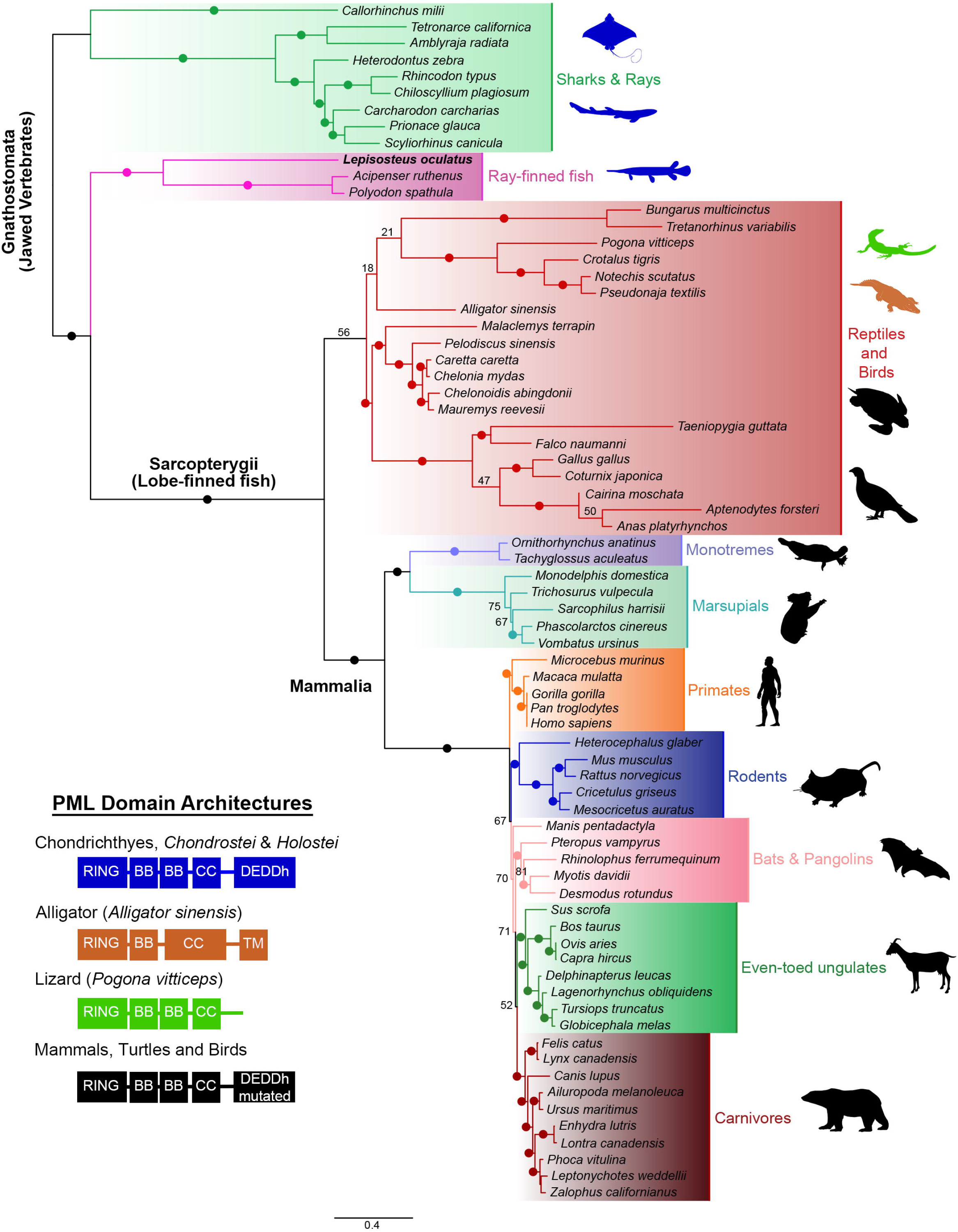
Domain architecture and Maximum Likelihood phylogenetic tree showing the relationships of PML homologs. 70+ protein sequences encoded in genomes from representative vertebrates were identified and utilized to construct the phylogenetic tree. Teleost fishes and amphibians lack full-length PML-I. Ultrafast bootstrap values are shown as numbers if <85% and indicated with solid dots on branches if >85%. Branch lengths indicate the expected number of amino acids substitution per site. Domains for homologs were determined using InterPro (https://www.ebi.ac.uk/interpro/) and 4 different domain architectures (left) were seen across analyzed sequences. Domain abbreviations refer to RBCC (Ring-B-box-Coiled Coil), BB (B-box), CC (Coiled coil), and DEDDh (3’-5’ DEDDh exonuclease). Sequences from species that were analyzed are shown in silhouettes and coloured according to the corresponding domain architecture. Silhouettes for species were obtained from PhyloPic (http://phylopic.org).

### ExoIII domain phylogeny

The phylogenetic analysis of TREX, Plex9, and PML-CDE domains (Figure 3B) shows that within metazoans, TREX and Plex proteins are distantly related and diverged before the last common ancestor of this group. Furthermore, this phylogeny shows that the ExoIII domains of PML are more closely related to Plex exonucleases. Syntenic analysis from the sequenced genomes revealed that TREX1 first appears in tetrapods due to its presence in amphibian genomes and that it likely arose from a gene duplication of TREX2. In contrast, genes encoding Plex9 sequences share synteny to the PML locus. Overall, this analysis is well supported by bootstrap values, highlighting the robust nature of the relationships uncovered, both between and within these two distinct exonuclease families.

### Plasmid construction

Synthetic genes coding for zebrafish Plex9.1 (Ile44-Leu244; Ensembl:ENSDARG00000092584; NCBI: XP_005174272.1) and spotted gar PML-CDE (Ala497–Glu766; Ensembl: ENSLOCG00000014935; NCBI: XP_015199146.1) were purchased (BioBasic Inc) and ligated into a modified pET21 expression vector that contained upstream sequences coding for a hexahistidine tag (H), maltose binding protein (M), and tobacco etch virus protease recognition sequence (T) to create pHMT-zfPlex9 and pHMT-sgPml-CDE, respectively. Site directed mutagenesis was used to create pHMT-zfPlex9.1 (D61N) using pHMT-zfPlex9.1 as a template. The fidelity of all plasmids was verified by sanger sequencing (Eurofins Genomics).

In addition, along with *zfplex9*.*1* and *sgpml*, another zebrafish gene coding for *plex9*.*2* (Ensembl ID: ENSDARG00000093773; NCBI: XP_002666829.1) was utilized in the study. In addition, axolotl TREX1/2/3 sequences (AMEXTC_0340000192613, AMEXTC_0340000056847, AMEXTC_0340000061536) from the axolotl transcriptome Am_3.4 available on Axolotl-omics (https://www.axolotl-omics.org). These coding gene sequences were all cloned into CMV-based FLAG-tagging and Clover-tagging vectors (CMV-FLAG-J1 and CMV-Clover-J1; derived from commercially available vectors pEGFP-C1 and pEGFP-N1; Clontech) for ectopic expression in human cells.

RNA used for cloning the axolotl *TREX*, spotted gar *pml*, and zebrafish *plex9*.*1* and *plex9*.*2* sequences was obtained from animal tissues (described below under animal ethics). Two gar individuals were raised as described in Darnet et al. (2019) to a total length of ∼30cm, euthanized with a lethal dose of MS-222 (Sigma); then tissues (muscle, gills, intestine, liver, brain, spleen, heart, gas bladder) were dissected, and stored in RNAlater (Ambion) (49). A caudal fin clip was stored in 95% ethanol. From the available tissue, expression of *sgpml* was highest in the spleen and thus this was utilized for downstream cloning. RNA for downstream cloning was derived from axolotl limbs that were dissected and stored in Trizol. In the case of zebrafish, the RNA is derived from 24 hpf zebrafish embryos. Quail RNA for quail PML cDNA was isolated from a pellet of QM5 cells (quail muscle 5 cells). The RNeasy (Qiagen) kit was used for all extractions, according to the manufacturer’s instructions. The cDNAs were prepared using the SuperScript™ IV One-Step RT-PCR System (Invitrogen) and sequenced prior to subcloning into indicated vectors.

For *Polyodon spathula* (NCBI: XP_041075354) and *Terrapene carolina triunguis* (Ensembl ID: ENSTMTG00000003177, NCBI: XP_024070906.1) *PML* coding gene sequence DNA was synthesized through gBlocks™ Gene Fragments in two parts (Integrated DNA Technologies). Overlap extension PCR was utilized to generate the full sequence, which was then cloned into the CMV driven FLAG-tag vectors used for the other coding sequences.

### Protein expression and purification of zfPlex9.1 and sgPml-CDE

BL21(DE3) *E. coli* (New England BioLabs Inc.) were transformed with pHMT-zfPlex9.1, pHMT-zfPlex9.1 (D61N) or pHMT-sgPml-CDE. Transformed cells were grown in LB media to an OD_600_ of ∼0.6, after which protein expression was induced with 0.5 mM IPTG. After overnight expression at 20 °C (16 °C for pHMT-zfPlex9.1 (D16N)), cells were resuspended in lysis buffer (HMT-ZfPLEX9.1: 20 mM Tris pH 7.5, 100 mM NaCl, 5 mM β-mercaptoethanol (BME); HMT-ZfPLEX9.1 (D61N): 20 mM CHES pH 9, 100 mM NaCl, 5 mM BME, 5 mM MgCl_2_; HMT-sgPml-CDE: 20 mM Tris pH 8, 500 mM NaCl, 5 mM BME), lysed by sonication, and clarified by centrifugation (25,000 × g for 20 min at 4 °C). The supernatant was loaded onto an amylose chromatography column (New England BioLabs Inc.), washed with lysis buffer, and eluted with lysis buffer containing 10 mM maltose. The proteins were then incubated with TEV protease overnight in elution buffer (HMT-ZfPLEX9.1) or with concurrent overnight dialysis at 4 °C against dialysis buffer (HMT-ZfPLEX9.1 (D61N): 20 mM CHES pH 9, 100 mM NaCl, 5 mM BME, 5 mM MgCl_2_; HMT-sgPml-CDE: 20 mM Tris pH 8, 250 mM NaCl, 5 mM BME). After cleavage, exonuclease domains were isolated using Ni^2+^ affinity chromatography and then purified by size exclusion chromatography (HiLoadTM 16/600 SuperdexTM 75 pg) using 20 mM Tris pH 7.5, 100 mM NaCl, 5 mM BME (HMT-ZfPLEX9.1 and HMT-ZfPLEX9.1 (D61N)) or diluted 5-fold with 20 mM MES (2-(N-morpholino) ethanesulfonic acid) pH 6, 5 mM BME and purified by cation exchange chromatography (HiTrap™ SP HP, cytiva). Purifications were monitored by SDS-PAGE and UV/Vis absorbance (using predicted extinction coefficients at 280 nm of 20970, 20970, and 18450 M^-1^cm^-1^ for ZfPLEX9.1, ZfPLEX9.1 (D61N), and sgPml-CDE, respectively).

### In vitro exonuclease assays

Unless otherwise noted, exonuclease reactions (40 µL) contained 500 nM 5’-6-FAM labeled oligonucleotide (ssDNA: 5’-ATACGACGGTGACAGTGTTGTCAGACAGGT-3’or dsDNA pseudo-palindrome: 5’-TCACGTGCTGAC/GTCAGCACGACG-3’), 20 mM Tris pH 7.5, 2 mM dithiothreitol, and 100 μg/mL BSA. For metal specificity assays the reactions contained 625 nM of zfPlex9.1 or sgPml-CDE and 2 mM metal (MgCl_2_, MnCl_2_, ZnSO_4_, or CaCl_2_) or 10 mM EDTA. Exonuclease titration assays contained 4.9–625 nM of zfPlex9.1 or zfPlex9.1 (D61N) and 5 mM MgCl_2_. Reactions were allowed to proceed for 20 min at room temperature and then quenched with 3 volumes of 100% ethanol, dried via speed vac, resuspended in 10 μL formamide, resolved by urea PAGE (19% 29:1 acrylamide and 7 M urea in TBE), and visualized using a fluorescence imager (VersaDoc).

### Microscale thermophoresis (MST) binding assays

Binding assays were carried out in 20 mM Tris pH 7.5, 100 mM NaCl, 5 mM CaCl_2_, and 5 mM BME. Solutions contained 50 nM ssDNA or dsDNA and zfPlex9.1, zfPlex9.1 (D61N), or sgPml-CDE to maximum concentrations of 11.7 µM, 15.2 µM, and 11.1 µM, respectively. Thermophoresis measurements were collected at 25°C using a Monolith NT.115 (NanoTemper Technologies) set to medium LED power and 20% excitation power. Raw data were processed using NT Analysis software (NanoTemper Technologies) and fit to a Hill binding model to determine the EC_50_ and Hill coefficient.

### Cell culture

U2OS osteosarcoma cell lines (parental U2OS, U2OS^Clover–PML^ (50), U2OS^GFP–PML-I^, U2OS TREX1 KO cells, U2OS PML KO cells (13)) were cultured in Dulbecco’s modified Eagle’s medium (Life Technologies) supplemented with 10% fetal calf serum, at 37°C with 5% CO_2_. QM5 (Quail Muscle clone 5) cells were a gift from the Roy Duncan lab (Dalhousie University) and grown like U2OS. NHDF (WT and PML KO cells) were cultured in alpha-modified Minimum Essential Medium (Life Technologies) supplemented with 15% fetal calf serum and GlutaMAX (Thermo Fisher) at 37°C with 5% CO_2_.

### Generation of a CRISPR/Cas9 KO lines

PML CRISPR/Cas9 KO lines, both immortalized NHDF and U2OS, were previously generated and characterized (13). We generated U2OS cells lacking *TREX1* expression, we designed a strategy to knock-in a puromycin resistance cassette into exon 2 of *TREX1*, thereby disrupting *TREX1* gene expression but not overlapping with the *ATRIP* protein encoding region. U2OS was used a cell line since they have impaired cGAS-STING signalling and allow for us to focus on L1-retrotransposition in an interferon-independent manner(51).

Two guide RNAs targeting the Cas9D10A nuclease to the *TREX1* locus were designed utilizing the CHOPCHOP v3 webtool for designing guide RNAs(52). We selected the top two guide RNAs, g1 (5’- CCCAACCATGGGCTCGCAGGCCCTGCCCCCGGGGCCCATGCAGACCCTCATCTTTTTCGACATGG-3’) and g2 (5’- CCATGTATGGGGTCACAGCCTCTGCTAGGACCAAGCCAAGACCATCTGCTGTCACAACCACTGCACACCTGG-3’) and cloned these guides into the All-in-one (AIO)-PURO vector encoding the Cas9D10A nickase (Addgene #74630). U2OS cells were treated with 2 µg/mL of puromycin (Invitrogen) and resultant clones were isolated. Resultant clones were screened by both western blotting and immunofluorescence to confirm that TREX1 expression was absent (Supplementary Figure S6).

We also examined *ATRIP* expression in the *TREX1* KO lines, which was previously ignored in other reports of a human *TREX1* KO line, despite *TREX1* being nested in the locus of *ATRIP (53,54)*. We were unable to get a positive clone with guide 2 (g2) and this appears to be a result of ATRIP levels being altered, where two prominent bands are present (data not shown). We therefore utilized the clone generated from guide 1 (which still show a faint secondary ATRIP protein band that is not present in WT cells) (Supplementary Figure S6). We utilized this clone as TREX KO#1 for experiments involving L1 suppression, a previously established function for TREX1 (both via KO models in mice and overexpression studies) (55,56). Our addback of TREX1 restored L1 suppression, supporting the notion that elevated L1 retrotransposition was a result of TREX1 loss and not ATRIP being altered (Figure 5).

### LINE retrotransposition assays

To assess the effect of proteins on retrotransposition, we utilized methods for previously characterized plasmids encoding a LINE retroelement (human L1, zfL2-1, zfL2-2 or UnaL1) that encode a neomycin cassette that when transfected into cells, is only successfully re-integrated upon retrotransposition (32,35,57-60). Upon G418 (Thermo Fisher) selection, it is possible to quantify the number of positive clones that survived which will be proportional to the LINE activity occurring. Co-expression of the proteins of interest with the LINE encoding plasmid, will determine which proteins enhance, suppress, or have no impact on cell LINE retrotransposition. HeLa and U2OS cells were utilized for LINE retrotransposition since they are susceptible to high levels of LINE retrotransposition in culture (32,57).

Cells (HeLa or U2OS) were seeded in a 6-well at densities of 2 × 10^5^ per well, left to attach for 24 hours and co-transfected using Lipofectamine 3000 (Roche) with (1) 1 ug of a retrotransposon encoding plasmid (2.5ug for the zfL2-1 retrotransposon) and (2) with 500 ng of a FLAG-tagged protein of interest. The only exception was the GFP-TREX1(D18N) plasmid which was obtained from Addgene (27220). No difference was observed between cell toxicity or transfection efficiencies between GFP-TREX1(D18N) and the other plasmids (data not shown). After 2 days post-transfection, the cells were reseeded on a 10 cm plate, allowed to adhere for 24 hours, and then incubated with G418 (500 ug/mL for HeLa; 400 ug/mL for U2OS) for a total of 10 days for HeLa and 14 days for U2OS. Then post-selection, the resistant colonies were fixed with 1 mL of methanol for 20 minutes. The colonies were then washed with PBS and stained with 0.5% crystal violet (stained for 30 minutes in 5% acetic acid and 2.5% isopropanol) for 30 minutes. The entire fixation and staining procedure were completed at room temperature. Colonies were then counted and retrotransposition activity was determined.

### 2’,3’-cGAMP quantification

4 × 10^6^ or 6 × 10^6^ U2OS cells (for TREX1 KO and PML KO experiments, respectively) were seeded into 15-cm dishes, and 24 hours later cells were transfected with the Bluescript vector (empty vector control) and Human L1 plasmid (used in L1 retrotransposition assay) using Lipofectamine 2000 reagent (Invitrogen). Cells were harvested 36 hours after transfection, washed with 2x with PBS and pelleted before lysis. Samples were resuspended in 500 μL M-PER (mammalian protein extraction reagent) lysis buffer (Thermo Scientific). Lysates were incubated on ice for 30 minutes with gentle agitation every 10 minutes, before being spun down at 16,000 x g, 4° C for 10 min. Samples were quantified using the 2′3′cGAMP ELISA Kit (Cayman Chemical) according to the manufacturer’s instructions.

### Immunofluorescence microscopy

For transfections, cells were seeded into wells containing coverslips in 6-well dishes, then transfected the next day with expression vectors. One day after transfection, coverslips were washed briefly with PBS and the cells were fixed in 2% PFA, permeabilized with 0.5% Triton X-100 in PBS and blocked with 4% BSA in PBS. Cells were then immunolabeled with primary antibodies specific for FLAG (mouse anti-FLAG M2, Sigma, F3165, 1:200), T7 (rabbit anti-T7, Millipore/Sigma, AB3790, 1:200), PML (two antibodies used; sheep anti-PML, Diagnostics Scotland, PML2A, 1:500; rabbit anti-PML, Bethyl Laboratories, A301-167A, 1:1000), SUMO (two antibodies used; mouse anti-SUMO-1, clone 21C7, Zymed, #33-2400, 1:200; rabbit anti-SUMO-1, Abcam, ab32058, 1:200), rabbit anti-DAXX (polyclonal D7810; Sigma-Aldrich, 1:500), rabbit anti-SP100 (Chemicon, 1380) and TREX1 (rabbit anti-TREX1, Abcam, ab185228, 1:400). Then for secondary staining, coverslips were washed with PBS and incubated with Alexa-Fluor 647 donkey anti-rabbit, Alexa-Fluor 488 donkey anti-sheep, Alexa-Fluor 488 donkey anti-rabbit, and Alexa-Fluor 555 donkey anti-mouse (Thermo Fisher Scientific) secondary antibodies. Finally, the cells were washed several times in PBS and incubated with 1 µg/mL of 4’,6-diamidino-2-phenylindole (DAPI) (Sigma) to visualize the nuclei.

For transfections, we marked ER and L1 RNP structures using plasmids obtained from Addgene and individuals. The ER was marked using a Cytochrome p450 (CytERM) tagged to mScarlet (Addgene #85066), which was utilized as an ER marker. T7-tagged ORF1p was previously used to mark L1 RNPs and encoded in the pES2TE1 vector. Plasmids for flag-tagging of proteins of interest are described above under plasmid construction, aside from FLAG-TREX1 (Addgene #27218), GFP-TREX1 (Addgene #27219) and GFP-TREX1 (D18N) (Addgene #27220) which were obtained from Addgene. U2OS cells were transfected using Lipofectamine 2000 (Invitrogen) and HeLa cells using Lipofectamine 3000 (Invitrogen), according to the manufacturer instructions.

Fluorescent micrographs were captured with a HQ2 charge-coupled device (CCD) camera (Photometrics) on a custom-built Zeiss Cell Observer Microscope (Intelligent Imaging Innovations) using a 1.3 NA 63X immersion oil objective lens and LED illumination via a Spectra light engine (Lumencor). Images were processed and analyzed using Slidebook (Intelligent Imaging Innovations) and Adobe Photoshop CS5.

### Western blotting

For western blot analysis, cells were recovered from confluent 10 cm culture dishes and washed with PBS. The cells were lysed on ice for 20 minutes in RIPA buffer (Sigma) with protease inhibitors (P8340, Sigma). Lysates were cleared (10 min, 15 000 × g, 4°C) and protein extracts were analyzed by SDS-PAGE and western blotting using 5% milk powder with 0.1% Tween 20 in PBS as a blocking solution. Protein was transferred to Nitrocellulose membrane (BioRad). Total protein was determined directly on the membrane using the 4-15% Mini-PROTEAN TGX-Stain Free Protein Gel system (BioRad). Antibodies used for Western blotting analysis were: rabbit anti-TREX1 (Abcam, ab185228, 1:1000) and anti-ATRIP (Sigma, PLA1030, 1:1000).

### Statistical analysis

All statistical analyses were performed using GraphPad Prism 9.0.1. The sample size and error bars for each experiment are defined in the figure legends. Comparisons between groups for LINE1 retrotransposition and the 2’3’-cGAMP assay were analysed by a repeated measures one-way ANOVA, with the Geisser-Greenhouse correction and then a Tukey’s multiple comparisons test between groups.

### Animal ethics

Spotted gar work was approved by the Institutional Animal Care and Use Committee at Michigan State University (protocol no. AUF 10/16-179-00). For axolotl work, Axolotl Université de Montreal animal ethics committee authorization was: 20-103. The Université de Montral animal ethics committee is recognized by the CCAC. The use of zebrafish embryos for generation of RNA was approved by Dalhousie University Committee on Laboratory Animals (Protocol Number 20-130)

## RESULTS

### PML first emerged in jawed vertebrates

To elucidate the evolutionary history of *PML*, we identified putative orthologs in publicly available databases of eukaryote genomes and transcriptomes (Figure 1). PML-I orthologs were identified in jawed vertebrates (Gnathostomata) including extant cartilaginous fishes (Chondrichthyes) and bony vertebrates (Euteleostomi) (61). We could not identify orthologs of PML in agnathan cyclostome (lamprey and hagfish) genomes (Figure 1). In addition, we documented the divergence of the *PML* gene in amniotes (Figure 1).

Among Chondrichthyes, PML ortholog architecture can be stratified into three groups, those with genomes encoding full length PML-I, those encoding putative orthologs with only a conserved RBCC domain, and those with a C-terminal transmembrane domain (Figure 1). While the RBCC domain is common among TRIM family proteins (62), the C-terminal domain of PML-I encoded by exon 9 uniquely shares homology with DEDDh exonucleases, which was originally annotated as a putative exonuclease III domain (42). This C-terminal DEDDh exonuclease (CDE) domain is conserved in bird, reptile, and mammalian PML orthologs as a predicted enzyme fold; however, the predicted catalytic residues are mutated in tetrapod orthologs of PML-I (Figure 1). Uniquely in ray-finned fish (such as paddlefish, spotted gar and sturgeon) and in cartilaginous fishes such as sharks and rays, PML orthologs have a CDE with intact catalytic residues, suggesting that they could be functional exonucleases (Figure 1).

During vertebrate evolution, the PML gene appears to have been lost several times in major extant euteleostome lineages, including amphibians and teleost fishes (Figure 1). We also failed to identify a PML ortholog in the coelacanth genome, the most basally diverging lineage of extant Sarcopterygii (lobe-finned fishes). Similarly, we could not identify an ortholog of PML in the lungfish genome. Within reptiles, turtles show conservation of the CDE, whereas lizard genomes encode a truncated PML ortholog lacking the CDE. Birds similarly have PML homologs that resemble full-length PML. Since turtles are thought to be more closely related to archosaurs than lepidosaurs, our analyses suggest that bird and turtle ancestral lineages retained an ortholog resembling full-length PML (63).

### PML was once a cytoplasmic exonuclease

The discovery of full length PML-I orthologs in ray-finned fish, sharks and rays encoding putatively active CDE domains was intriguing. We utilized the spotted gar genome to further explore PML-I from ray-finned fish (64). The spotted gar lineage diverged ∼350 million years ago from the teleost lineage before the teleost genome duplication event and has a slowly evolving genome (64). These features make the spotted gar an excellent representative species for studying the *PML* gene in fish. At the locus encoding the spotted gar *pml* ortholog (referred to as *sgpml*), there is strong synteny to the human *PML* locus on human chromosome 15, with flanking genes *STOML1, ISLR, CCDCC33* being conserved within the syntenic region (Figure 2A). We also utilized a bridging amniote species, *Coturnix japonica* (Japanese quail) that encodes a *PML* locus (referred to as *jqPML*) with synteny to both the spotted gar and human loci (Figure 2A).

**Figure 2.**
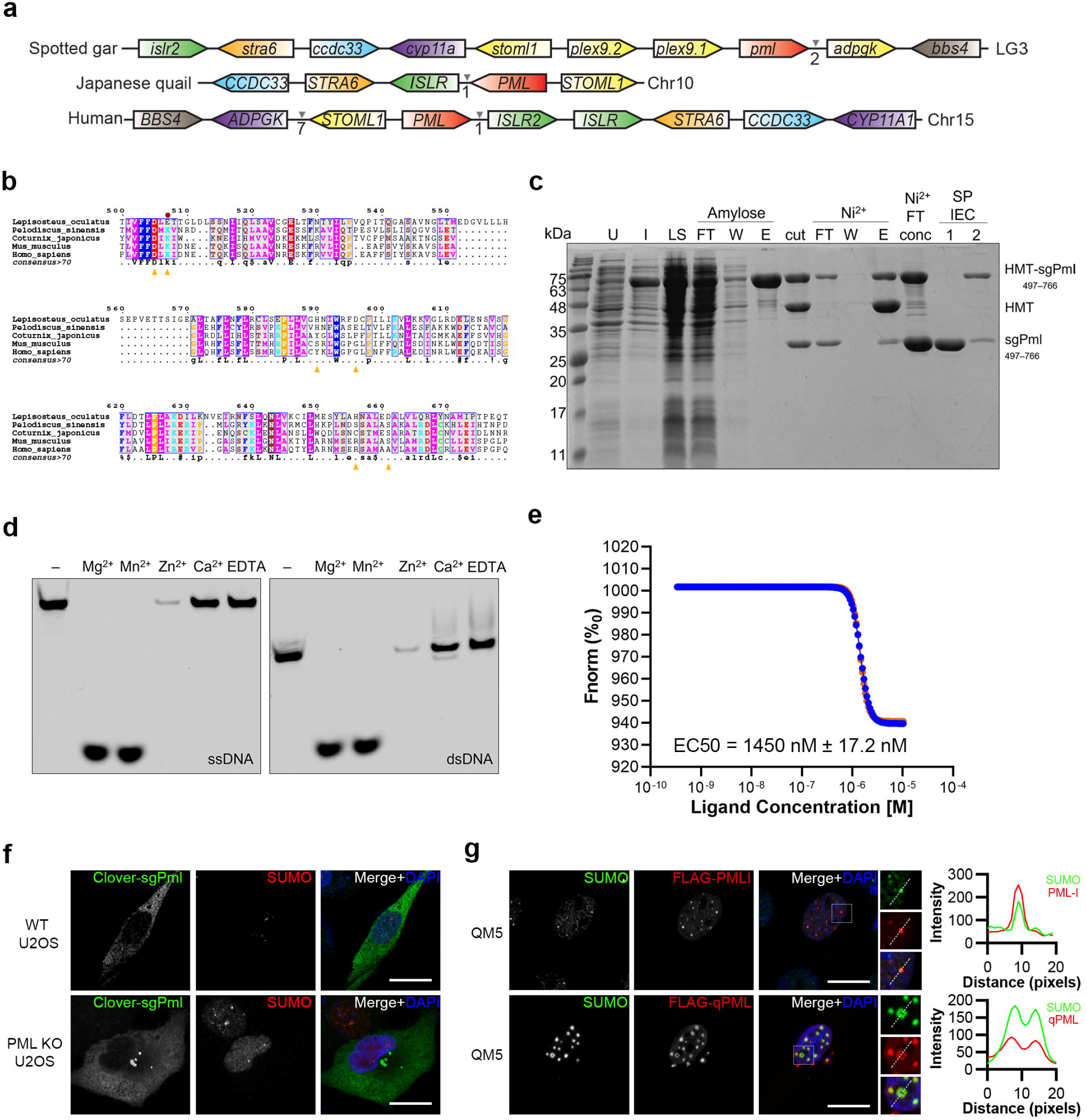
Spotted gar Pml is an active cytoplasmic exonuclease. (a) Synteny of *pml* locus between the genomes of the Spotted gar, Japanese quail and Humans. Similarly coloured genes represent homologs between species. The chromosome where the *pml* locus is found in each species is indicated on the right. The intervening arrows with numbers indicate the number of intervening adjacent genes that exist between the homologous genes not found in the syntenic region. (b) MUSCLE alignment of the C-terminal DEDDh exonuclease domain identified in the spotted gar PML (sgPml) protein with other vertebrate PMLs. The consensus sequence is displayed below the sequences for residues with over 70% conservation across species. Yellow arrows indicate predicated catalytic residues for sgPml and the red circle indicates the conserved SUMOylation site (K616 in humans) found in amniote species. (c) Purification of sgPml-CDE. sgPml-CDE was expressed as a fusion to sequences encoding for hexahistidine, maltose binding protein, and a TEV protease cleavage site (HMT). SDS-PAGE analysis is shown for samples of uninduced (U) and induced (I) cells, soluble lysate (LS), after TEV cleavage (cut), flowthrough (FT), wash (W), and elution (E) fractions of amylose and Ni2+ affinity chromatography, and fractions from ion exchange chromatography (IEC). (d) Exonuclease activity of sgPml-CDE requires Mg2+ or Mn2+. Fluorescent oligonucleotides were incubated with sgPml-CDE in the presence of the indicated divalent cation or EDTA. “-” refers to input without the addition of cations. (e) sgPml binds to dsDNA with high affinity. Microscale thermophoresis was used to quantify the affinity between sgPml-CDE and dsDNA (EC50 = 1450 nM ± 17.2 nM; n=2). The different coloured lines are data shown for two replicates. (f) sgPml localizes to the cytoplasm unlike human PML. Ectopic expression of Clover-tagged sgPml in U2OS cells shows that the protein localizes to the cytoplasm and forms SUMO-independent bodies. (g) Quail PML (qPML) localizes to both the nucleus and cytoplasm in quail cells. qPML is found SUMOylated in the nucleus akin to human PML. Scale bars represent 10 µM for (f) and (g).

Given that the sgPml protein encodes a potentially active CDE (Figure 2B), we cloned the full length *sgPml* gene from gar splenic tissue for further characterization. We also expressed the codon optimized putative DEDDh exonuclease of sgPml (residues 497-766) and purified it to homogeneity (Figure 2C). We found that the sgPml-CDE encodes an active exonuclease capable of degrading ssDNA and dsDNA when incubated with Mg^2+^ and Mn^2+^ (Figure 2D). Thus, akin to other DEDDh family exonucleases (65), it appears that sgPml can degrade dsDNA in the presence of divalent cations. In addition, using microscale thermophoresis binding assays in the presence of Ca^2+^, we found that sgPml-CDE can bind to dsDNA with an EC50 of 1450 nM ± 17.2 nM (Figure 2D) but bound ssDNA at lower affinity. These data indicate that sgPml has the capacity to bind and degrade both ssDNA and dsDNA but with a higher affinity for dsDNA through its CDE.

### Nuclear PML bodies emerged in amniotes

We next characterized the cellular localization of sgPml by expressing it in human wild type (WT) and PML KO U2OS osteosarcoma cells. Strikingly, in contrast to mammalian PML protein, sgPml did not form nuclear bodies and localized diffusely in the cytoplasm, forming ∼1-3 cytoplasmic body-like puncta (Figure 2F). We also expressed the American Paddlefish PML ortholog, another predicted PML exonuclease and observed cytoplasmic localization like sgPml (Supplementary Figure S1). Since mammalian PML NBs are associated with post-translation modifications of nuclear body proteins by the small ubiquitin like modifier (SUMO) (63), we also examined SUMO-1 localization relative to these cytoplasmic sgPml puncta. These sgPml puncta were negative for SUMO1 immuno-staining, and therefore form in a SUMO-independent manner, unlike mammalian PML NBs (Figure 2F).

Sequence comparisons revealed that the target lysines involved in human PML SUMOylation, which first appear in amniote species according to our evolutionary reconstruction, align partly to the CDE and its catalytic residues (Figure 2B). This could suggest that a shift in PML cellular function and localization may have occurred concurrently with the acquisition of SUMOylation. To assess the possibility that SUMO site evolution drove the nuclear localization and function of PML, we cloned and expressed the Japanese quail PML ortholog (jqPML) (Figure 2B). The expression of jqPML, human PML-I, and human PML-IV in quail cells all led to the formation of SUMO colocalizing nuclear bodies (Figure 2G). However, like sgPml, jqPML also formed cytoplasmic bodies lacking SUMO (Figure 2G). The expression of a PML ortholog from a turtle species mirrored the localization of jqPML (Supplementary Figure S1). jqPML NBs also co-localized with canonical PML NB proteins, DAXX and SP100 (66) (Supplementary Figure S1). However, turtle PML NBs only appeared to recruit DAXX and were void of SP100 (SP100 foci are neighboring). These results support the importance of PML SUMOylation for nuclear body formation, and indicates that avian and turtle PML orthologs, with the acquisition of SUMO-target lysine residues, can localize to both the cytoplasm and the nucleus to form PML NBs.

### Plex9 proteins are novel DEDDh exonucleases

Most ray-finned fish species (99.8%) are teleost fish that underwent a genome duplication event accompanied by subsequent gene order rearrangements (67). During this event, it appears that the full length *PML* gene was lost in teleost fishes. However, we identified a pair of genes sharing a high degree of homology to the CDE of sgPml in zebrafish. We describe these novel genes as <u>P</u>ml-<u>l</u>ike <u>ex</u>on <u>9</u> genes (*Plex9*.*1* and *Plex9*.*2*), which like the PML-CDE, share sequence similarity with TREX1 and TREX2, the two major DEDDh exonucleases in mammals (Supplementary Figure S2A) (65,68). A closer examination of the spotted gar *pml* locus also revealed the presence of two adjacent upstream genes that appear to have duplicated from the PML-CDE (Figure 3A). Phylogenetic analyses also indicated that Teleostei Plex9 shares the highest similarity to the PML-CDE, and across genera, TREX1 orthologs consistently cluster independently from Plex9 orthologs (Figure 3B). Thus, *plex9* genes in Telostei are DEDDh exonucleases closely related to the CDE of PML-I and are not evolutionarily related to *TREX1/2*.

**Figure 3.**
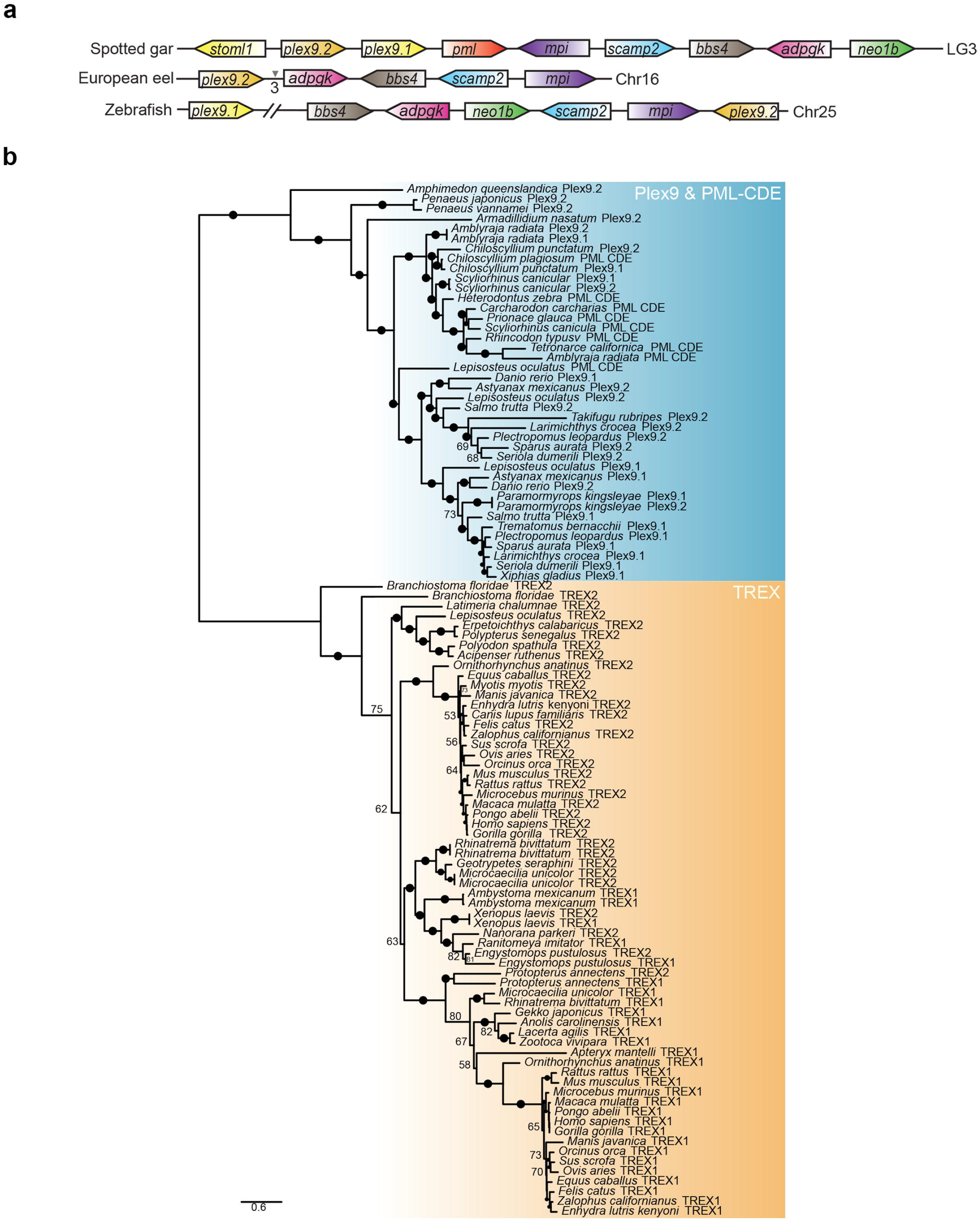
Origins of PML-like exon 9 (*Plex9*) and *TREX1* genes. (a) Teleost *plex9* genes show synteny to the spotted gar *pml* locus. Similarly coloured genes represent homologs between species. The chromosome where the *pml* locus is found in each species is indicated on the right. The intervening arrows with numbers indicate the number of intervening adjacent genes that exist between the homologous genes not found in the syntenic region. (b) Plex9 homologs resemble both TREX1/2 and PML exonucleases but are related to PML exonucleases. A maximum likelihood tree was constructed using Plex9, PML CDE and TREX1/2 protein sequences. While the TREX1 proteins cluster with TREX2, sequences for Plex9 cluster with PML CDE sequences. Bootstrap values are indicated beside each node. Ultrafast bootstrap values are shown as numbers if <85% and indicated with solid dots on branches if >85%. Branch lengths indicate the expected number of amino acid substitutions per site.

We also identified Plex9 orthologs in other ray-finned fishes and in the elasmobranch sub-class of cartilaginous fish, whereas the TREX1 gene first appears in the tetrapod ancestor since it is found in amphibian species and amniotes (Figure 3B). Neither *plex9*.*1* or *plex9*.*2* orthologs can be identified in lobe-finned fish (Figure 3B). TREX2 predates TREX1 and can be found conserved in cephalochordates such as *Branchiostoma floridae*. Syntenic analysis of the *TREX1* loci reveals that *TREX1* was likely derived from a duplication of the *TREX2* gene (Supplementary Figure S2B). Thus, while tetrapods lost the *Plex9* genes, they retained the newly emerged *TREX1* gene and the *PML* gene encoding a CDE domain. In amphibians, we identified a putative ortholog of *TREX1* but could not identify genes resembling *PML* or the *plex9* orthologs. The axolotl genome encodes a TREX1 ortholog and there is expression of three transcripts for TREX1-like proteins (*axTREX1, axTREX2, axTREX3*). Previously, it was thought that *TREX1* was exclusive to mammals (69), however it appears that the progenitor *TREX1* gene encoding a DEDDh exonuclease is more generally present in tetrapod genomes.

To better understand the function of the Plex9 and non-mammalian TREX1 proteins, we cloned the zebrafish *plex9* genes (*zfplex9*.*1* and *zfplex9*.*2*), as well as the TREX1 ortholog isoforms from the axolotl genome (*axTREX1, axTREX2, axTREX3*) to assess their cellular roles and localization in mammalian cell assays (Figure 4B). Expression of these orthologs in U2OS cells revealed that zfPlex9.1 as well as axTREX1 and axTREX2 localized to the ER and cytoplasm in a similar manner as TREX1 (Figure 4C), whereas zfPlex9.2 localized primarily to the cytoplasm (Figure 4C). Like TREX1 which has recently been linked to nucleotide excision repair (70-73), zfPlex9.1 translocated to the nucleus in response to DNA damage (Supplementary Figure S3A). Thus, these different DEDDh exonucleases localize similarly to human TREX1 (Figure 2B).

**Figure 4.**
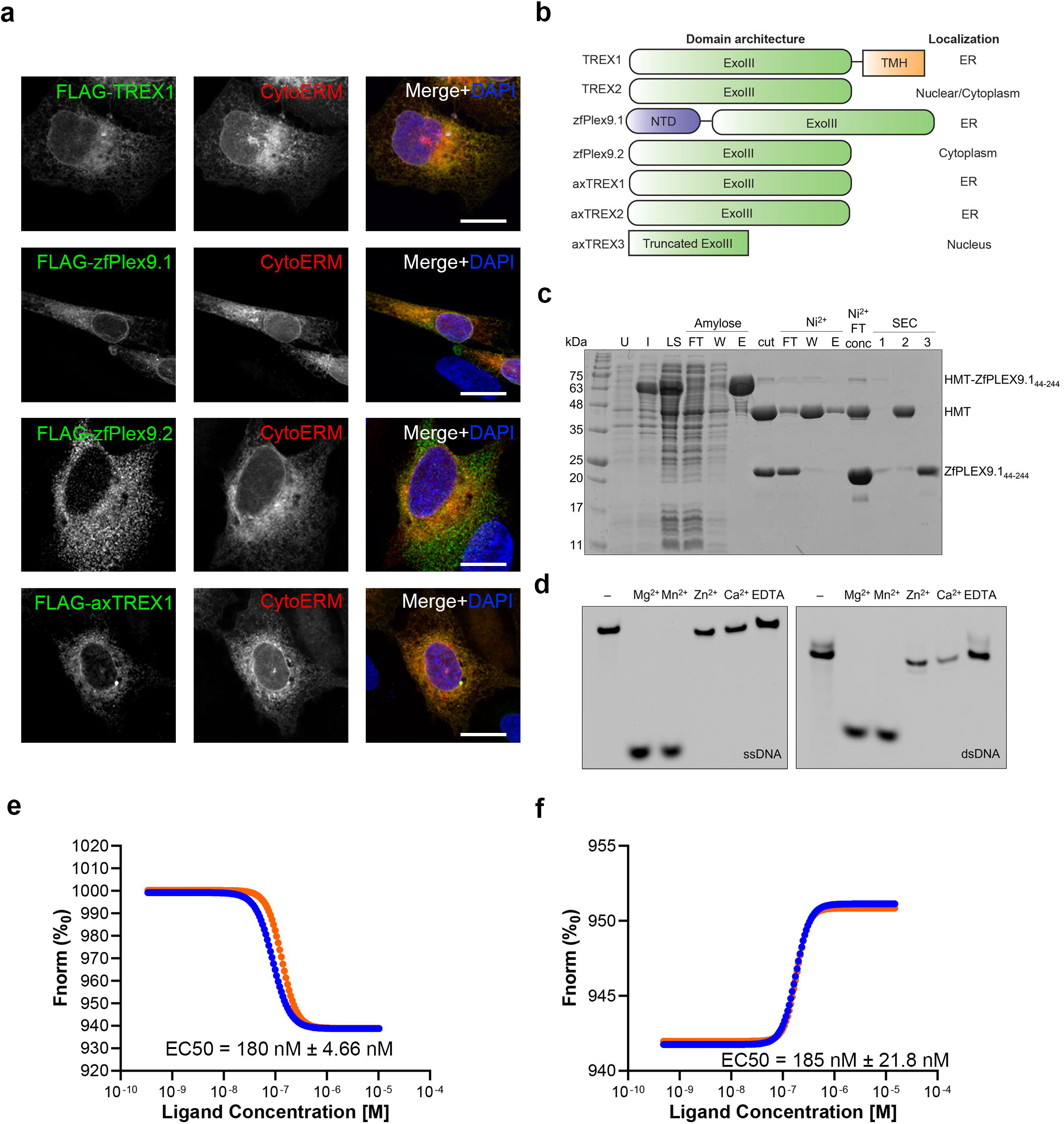
Plex9 proteins represent a novel class of DEDDh DNA exonucleases. (a) Ectopic expression of zebrafish Plex9.1/.2 (zfPlex9.1/.2) and axTREX1/2 reveals that these proteins localize to the ER and cytoplasm. Proteins were FLAG-tagged and expressed in U2OS cells to assess localization. A domain of Cytochrome p450 (CytERM) tagged to mScarlet (addgene #85066) was utilized as an ER marker. (b) The different domain architectures of TREX1/2 and Plex9 proteins and their known/determined localizations are shown on the right. TREX2 localization was assessed previously as both nuclear and cytoplasmic. (c) Purification of zfPlex9.1. zfPlex9.1 was expressed as a fusion to sequences encoding for hexahistidine, maltose binding protein, and a TEV protease cleavage site (HMT). SDS-PAGE analysis is shown for samples of uninduced (U) and induced (I) cells, soluble lysate (LS), after TEV cleavage (cut), flowthrough (FT), wash (W), and elution (E) fractions of amylose and Ni2+ affinity chromatography, and fractions from size exclusion chromatography (SEC). (d) and (e) zfPlex9.1 requires Mg2+ or Mn2+ for activity. Fluorescent oligonucleotides were incubated with zfPlex9 and the indicated metal. Samples were resolved by UREA-PAGE and visualized via fluorescence imaging. “-” refers to input without the addition of cations. (f) and (g) DNA binding of zfPlex9.1. Microscale thermophoresis was used to quantify the affinity between zfPlex9.1 with (f) ssDNA (EC50 = 185 nM ± 21.8 nM) and (g) dsDNA (EC50 = 180 nM ± 4.66 nM) (n=2). The different coloured lines are data shown for two replicates. Scale bars represent 10 µM for (a).

We next assessed the exonuclease activity of recombinant zfPlex9.1 (aa 44-244) and found it could degrade ssDNA and dsDNA (Figure 4C-E) in the presence of Mg^2+^ and Mn^2+^. It also bound to ssDNA (EC_50_ = 185 nM ± 21.8 nM) and dsDNA (EC_50_ = 180 nM ± 4.66 nM) with higher affinity than sgPml (Figure 4F). An alignment of zfPlex9.1, zfPlex9.2 and TREX1 revealed a conservation of the DEDDh catalytic residues (Supplementary Figure S2A) that allowed us to phenocopy the exonuclease deficient D18N TREX1 mutation by mutating the cognate residue in zfPlex9.1 (i.e., D61N; Supplementary Figure S2B). The D61N mutation impaired zfPlex9.1 catalytic activity without affecting the ability to bind both ssDNA and dsDNA (Supplementary Figure S4).

An important role for DNA exonuclease function for TREX1 is the clearance of micronuclei (74). To determine whether zfPlex9.1 and axTREX1 proteins were degrading micronuclei akin to TREX1, we treated U2OS cells with reversine, which induces micronuclei by causing the mis-segregation of chromosomes (56). We found that, just like TREX1, zfPlex9.1 and axTREX1 also accumulated at micronuclei (Supplementary Figure S3B). Thus, the Plex9 proteins encode active exonucleases with specificity towards ssDNA and dsDNA, which likely contributes to their ability to facilitate the clearance of micronuclei.

### sgPml and Plex9 proteins suppress L1 retrotransposition

TREX1 is the essential brake of the cGAS-STING pathway in mammals and has been shown to prevent genome instability by degrading cytosolic DNA and micronuclei, in addition to repressing endogenous LINE-1 (L1) retrotransposons (55,56,74). The cGAS-STING pathway is highly conserved in metazoans (75). Despite lacking a *TREX1* gene homolog, teleost fishes have a functional cGAS-STING pathway (76,77), raising the possibility that another DEDDh exonuclease may exist that functionally replaces TREX1. Given that teleost *plex9*.*1* and *plex9*.*2* genes encode active DEDDh exonucleases capable of degrading ssDNA and dsDNA that may be involved in micronuclei clearance, we hypothesized that these proteins may have functionally converged with TREX1 to compensate for its absence in the teleost lineage.

The role of TREX1 in L1 suppression is independent of catalytic exonuclease function (56). Similarly, the related SAMHD1 restriction factor, also suppresses L1 in a catalytically independent manner (78). Both TREX1 and SAMHD1 appear to restrict L1 through their nucleic acid binding functions (56,78). The Plex9 proteins and sgPml both bind nucleic acids and are active exonucleases. Given their mutual exclusivity with TREX1 in the fish genomes in which they are found, we next assessed if they functionally replace TREX1 in L1 suppression. To accomplish this, we employed the neomycin-resistance based clonogenic L1 assay that was previously used to demonstrate the role of TREX1 in regulating L1 (Supplementary Figure S5A) (74), to determine if these zfPlex9 exonucleases, axTREX1/2/3, and the related sgPml, could also suppress L1 retrotransposition. As reported previously, TREX1 and exonuclease deficient TREX1 (D18N) both significantly suppressed human L1 activity significantly (p<0.0001, Figure 5A). Consistent with a convergent role in the suppression of L1 retrotransposition, zfPlex9.1 and zfPlex9.2, axTREX1 and axTREX2, and sgPml all potently suppressed L1 activity (Figure 5A). Like TREX1 (D18N), exonuclease-deficient zfPlex9.1 (D61N) also suppressed L1 retrotransposition (Figure 5A).

**Figure 5.**
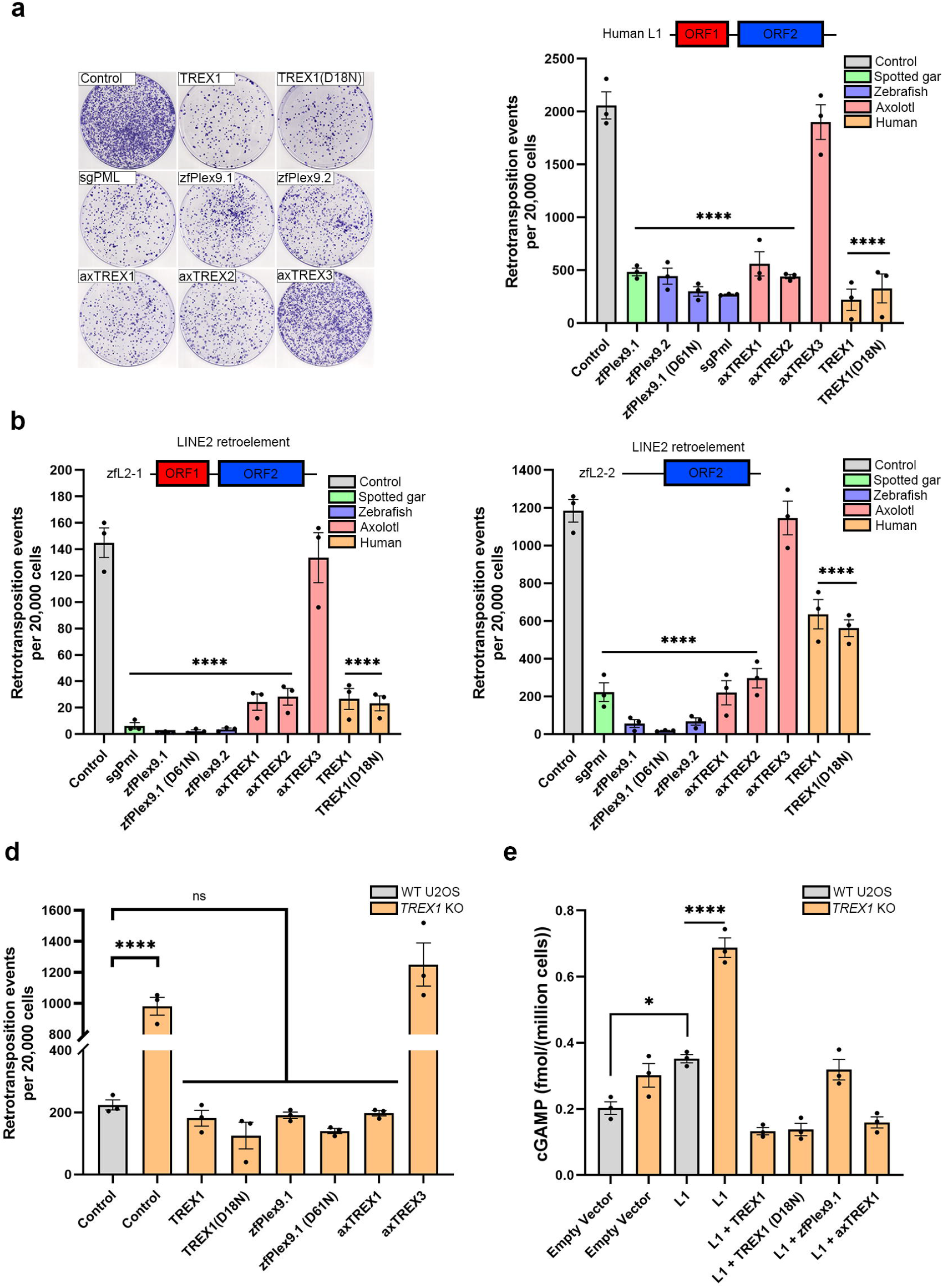
sgPml, Plex9 and axTREX1 proteins suppress LINE-1 (L1) activity in an exonuclease-independent manner. (a) Retrotransposition of human L1 was assessed with the co-expression of the FLAG-tagged versions of indicated proteins. Resultant plates from the assay (right) were stained with 0.5% crystal violet and quantified. Retrotransposition activity of zebrafish L1 elements were also assessed in (b, left) zfL2-1 and (b, right) zfL2-2 (n=3 for human L1, zfL2-1 and zfL2-2). (d) A *TREX1* knockout was generated using CRISPR/Cas9 and validated in U2OS (details in Supplementary Figure S4). (d) Retrotransposition activity was elevated in *TREX1* knockouts and addback of the zfPlex9.1 and zfPlex9.2, as well as axTREX1 and axTREX2 reversed this phenotype (n=3). (e) 2’,3’-cGAMP levels in U2OS WT and *TREX1* knockout cells after the transfection of an empty vector and the human L1 vector (n=3). 2’,3’-cGAMP concentrations were normalized to cell number for each experiment. Except for GFP-TREX1(D18N) which was previously obtained from addgene, the other proteins were FLAG-tagged. Controls represent empty vector (FLAG-tag backbone) used in the co-transfection with the L1 plasmid.

While L1 is active in mammals, related endogenous retroelements exist in other metazoan lineages. For example, the related LINE-2 (L2) elements are active in echinoderms and teleost fishes, but not mammals. Several teleost LINE elements have been characterized (79). We wanted to determine if zfPlex9 and sgPml proteins could suppress these active L2 elements. We again utilized neomycin-based retrotransposon reporters encoding teleost L2s (zfL2-1, zfL2-2 and UnaL2) (80). Ray-finned zfPlex9.1, zfPlex9.2, and sgPml potently suppressed the retrotransposition of both zfL2-1 and zfL2-2 (Figure 5B,C). Amphibian axTREX1 and axTREX2, also suppressed the teleost L2 elements like the Plex9 proteins (Figure 5B,C). However the truncated axTREX3 protein had no effect on L1 or L2 suppression (Figure 5B,C). We observed a similar relationship between the Plex9 proteins and axTREX1 with respect to UnaL2 (Supplementary Figure S5). Human TREX1, although capable of suppressing zfL2-1 and zfL2-2, was not as efficient at suppressing these L2 elements compared to the teleost Plex9 proteins, suggesting a degree of species tropism between DEDDh exonucleases and their target retroelements. Therefore, it appears that Plex9 and TREX1 orthologs broadly inhibit LINE retroelements, including L2 elements found in other vertebrate species.

We next sought to determine if the Plex9 proteins could functionally complement the increase in L1 activity previously demonstrated for TREX1 loss in human cells (56,74). Thus, we generated TREX1 knockout (KO) U2OS lines (Supplementary Figure S6). As expected, TREX1 KO led to a significant increase in human L1 retrotransposition, which was reversed by the overexpression of FLAG-TREX1 or GFP-TREX1(D18N) (Figure 5D). Consistent with being true TREX1 orthologs in axolotl, the axTREX1 and axTREX2 proteins encoding full-length DEDDh domains also successfully inhibited L1 retrotransposition when over expressed in TREX1 KO U2OS cells (Figure 5D). The ectopic expression of zfPlex9.1 and zfPlex9.2 also significantly reduced human L1 retrotransposition in the TREX1 KO cell line (Figure 5D). Thus, zfPlex9.1 and zfPlex9.2 have a similar convergent function to TREX1 with respect to suppression of LINE retroelements. TREX1 inhibits the production of 2’,3’-cGAMP by CGAS after L1 activity occurs (81). We found that the significant increase in 2’,3’-cGAMP caused by TREX1 loss with the transfection of the L1 plasmid could be reversed by the addback of zfPlex9.1 and axTREX1 (Figure 5E). Together, these results are consistent with an early role for TREX1 in the suppression of L1 activity in amniotes, and that in species that lack TREX1 such as zebrafish and other ray-finned fish, Plex9 proteins converged in function to similarly suppress L1 retroelements and downstream cGAS-STING signalling.

### Human PML-I shuttles to the cytoplasm to restrict L1 activity

While human PML retained the DEDDh exonuclease fold, the catalytic residues are changed relative to sgPml, and there is no evidence to suggest it is an active exonuclease (Figure 2B). To date no cellular function has been ascribed to the PML-I-CDE (Figure 6A). Since we and others have shown that L1-suppression is an exonuclease-independent function for TREX1 and the Plex9 proteins (Figure 5), we hypothesized that human PML may have retained a role in suppressing L1 activity via the CDE.

**Figure 6.**
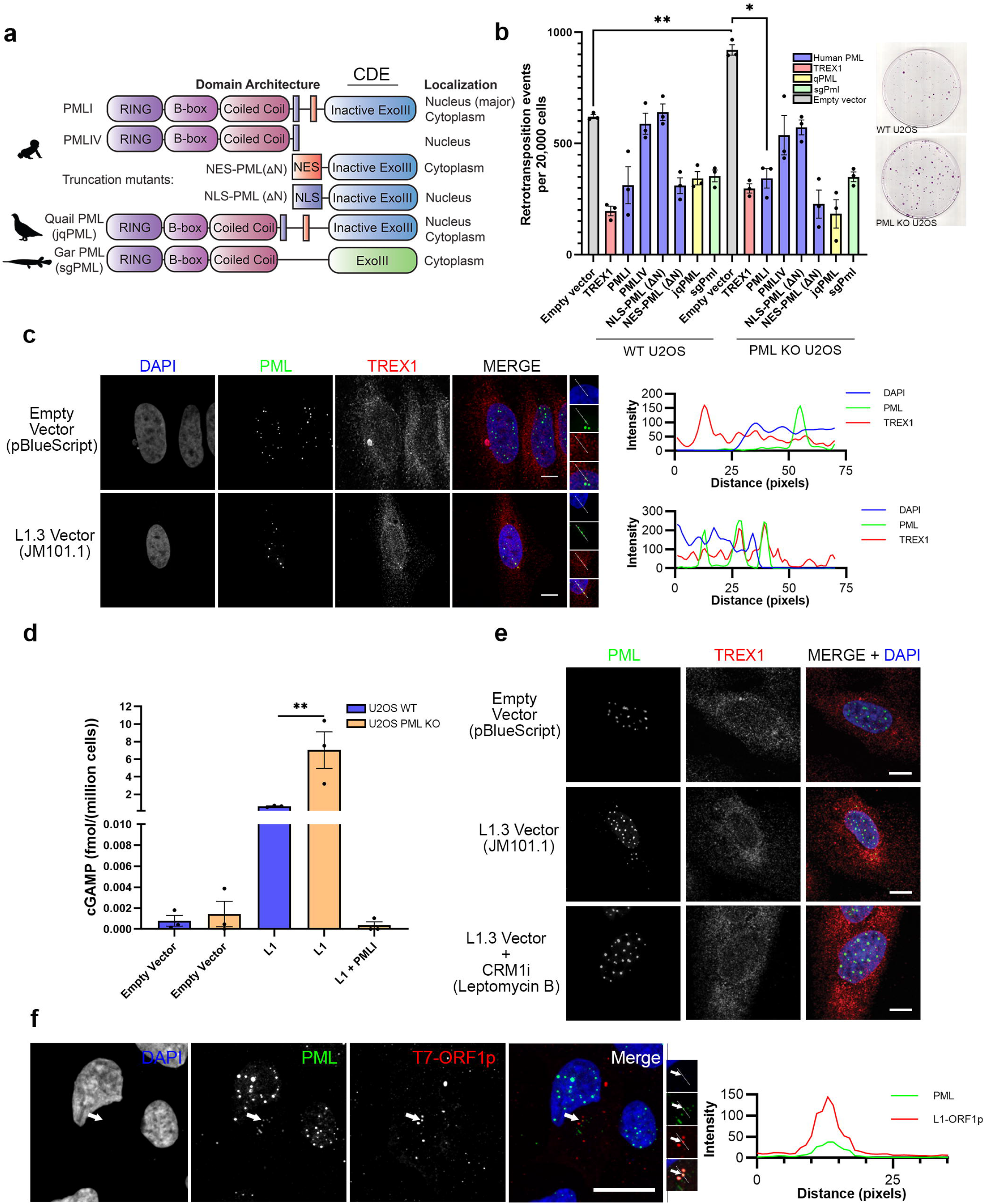
Human PML shuttles to the cytoplasm suppress L1 activity through its C-terminal DEDDh exonuclease (CDE) domain. (a) PML mutants were generated encompassing the CDE (aa positions 596-882) and tagged with both an SV40 NLS and the PML-I NES at the N-terminus. Localizations of the mutants are shown in Supplementary Figure S6. The mutants lack the C-terminal RBCC domain of PML. (b) Loss of *PML* elevates L1 activity which can be reversed with the addback of PML-I and the NES-PML mutant (n=3). Resultant plates from the assay were stained with 0.5% crystal violet and quantified. (c) Transfection of the human L1 retrotransposition vector led to cytoplasmic PML puncta forming. PML co-localized partly with TREX1 in the cytoplasm after L1 retroelements were active. (d) 2’,3’-cGAMP levels in U2OS WT and *PML* knockout cells after the transfection of an empty vector and the human L1 vector (n=3). 2’,3’-cGAMP concentrations were normalized to cell number for each experiment (e) The nucleocytoplasmic shuttling of PML-I is CRM1-dependent. Treatment of U2OS cells overexpressing GFP-PML-I for 3 hours with 10 ng/mL of Leptomycin B, a potent inhibitor of CRM1, a nuclear export protein, led to retention of GFP-PML-I in the cytoplasm. (f) Cytoplasmic PML partly localizes to ORF1p puncta in the cytoplasm. U2OS WT cells were transfected with human L1 encoding T7-tagged ORF1p. Scale bars represent 10 µm for images in (c), (e) and (f).

To test this hypothesis, we co-transfected HeLa cells with the human L1 reporter with FLAG-tagged human PML-I, PML-IV or jqPML; the latter to demonstrate conserved PML function in L1 suppression in other amniotes. PML-IV is almost identical to PML-I but lacks the CDE domain encoded by exon 9 (aa positions 596-882) (Figure 6A). We found that PML-I, but not PML-IV, significantly suppressed L1 activity (p<0.05) (Supplementary Figure S7). We next determined if the loss of endogenous PML influenced L1 retrotransposition in PML KO U2OS cells(13). KO of PML in U2OS cells increased L1 activity nearly 2-fold (p<0.01) compared to WT cells, which was reversed by expression of PML-I, jqPML or sgPml (Figure 6B). Collectively these data indicate that PML orthologs from 3 distinct vertebrate species, quail, human and gar, are capable of robustly suppressing human L1 activity.

In addition to encoding a C-terminal DEDDh exonuclease-like domain, PML-I also uniquely encodes a nuclear export signal (NES) for which no known stimulus has been identified to trigger nucleocytoplasmic shuttling (43). We also observed PML colocalize with Clover-sgPml in the cytoplasm in a small population of stressed cells (Supplementary Figure S8A). Therefore, we sought to determine if PML nucleocytoplasmic shuttling occurs directly in response to the stress of L1 retrotransposition. Indeed, in cells transfected with the active human L1 retroelement, endogenous PML shuttled robustly to the cytoplasm (Figure 6C). In the cytoplasm, PML formed large SUMO-negative puncta that resembled cytoplasmic sgPml bodies, which also co-localized with TREX1 puncta (Figure 6C). Like in the absence of TREX1, we observed that 2’,3’-cGAMP levels were elevated significantly (p<0.01) in the absence of PML after L1 transfection (Figure 6D). The increase in 2’,3’-cGAMP could be suppressed by the addback of PML-I into the PML KO cells (Figure 6D).

Since PML-I encodes a putative NES (positions 704–713) and has been identified as an interactor of CRM1 (63,82), we next examined if PML shuttling required CRM1/XPO1, a major nuclear export protein involved in NES-dependent shuttling. To determine if PML-I shuttling was dependent on CRM1, we utilized the CRM1-inhibitor leptomycin B (83). In the presence of leptomycin B, we found that PML-I shuttling to the cytoplasm did not occur (Figure 6C). Together, these data indicate that PML nucleocytoplasmic shuttling occurs in a CRM1-dependent manner in response to active L1 retrotransposition.

Substantial PML protein is still retained in the nucleus even under conditions of high L1 activity, it raises the possibility of a nuclear function for the DEDDh domain of PML in suppressing L1 retrotransposition. To determine if this is the case, we generated N-terminus truncation mutants of PML that encoded the PML-I C-terminus (aa 597-882) with either the upstream putative NES or a nuclear localization signal from SV40 (SV40-NLS) (Supplementary Figure S8B). We observed that expression of the NES-PML mutant potently suppressed L1 activity, in a manner comparable to jqPML and sgPml (Figure 6B,C). In contrast, the constitutively nuclear NLS-PML-I C-terminus had no significant effect on L1 retrotransposition (Figure 6B). Cytoplasmic PML also localized to ORF1p-positive structures, indicating a direct role for PML-I in the suppression of L1 Ribonucleoproteins (Figure 6E). These results collectively indicate an evolutionary conserved role for PML in suppressing L1 elements, which it accomplishes by CRM1-dependent nucleocytoplasmic shuttling of PML-I and requires its NES and C-terminal DEDDh domain.

## DISCUSSION

This study of PML gene evolution highlights how the availability of newly sequenced genomes from diverse taxa can help illuminate the complex molecular evolution of vertebrate genes and the functions they encode. Here, we determined that PML-I encodes a vestigial DEDDh exonuclease domain that has a novel evolutionarily conserved function in L1 suppression in humans and other amniotes. The PML protein has changed significantly over the course of jawed vertebrate evolution, both in cellular localization and function, which seems to be intricately tied to its SUMOylation. For example, human PML-I is primarily a nuclear protein acting as a scaffold for PML NB formation, while gar and paddlefish PML localize to the cytoplasm (Figure 7). Nonetheless, human PML-I can still facilitate L1 suppression through the acquired ability to shuttle to the cytoplasm via its nuclear export signal. This feature represents a remarkable compensation mechanism that allows PML to retain its key role in surveillance and suppression of L1 retroelement activity despite its divergence during amniote evolution, which is consistent with the broader role of PML in antiviral innate immune pathways (3,4).

**Figure 7.**
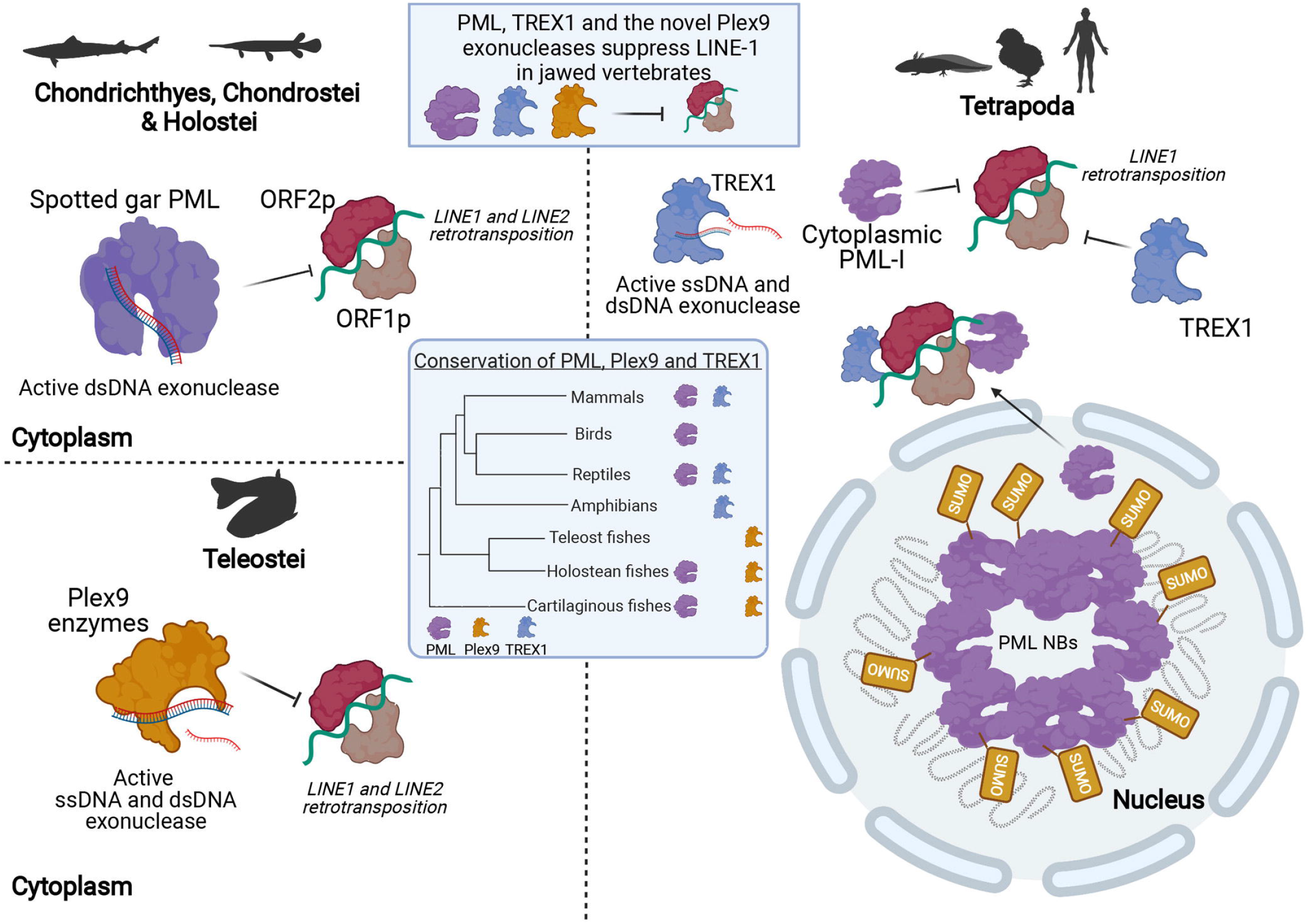
Overview of the suppression of LINE-1 retrotransposition in jawed vertebrates through the collective functions of PML, Plex9 and TREX1 enzymes. In fish, PML and novel Plex9 enzymes play a major role in suppressing both LINE-1 and LINE-2 elements. Despite the loss of *PML* in Teleost fish genomes, they have retained *Plex9* genes for the purpose of restricting LINE retrotransposition. *TREX1* appears later in tetrapods, where *Plex9* genes are lost. In addition, enzymatic PML function is lost and the protein forms nuclear bodies rather than localizing to the cytoplasm. However, PML has a retained an exonuclease-independent role for the surveillance of LINEs by shuttling to the cytoplasm to suppress the LINE1 retrotransposition. The three families of protein (PML, Plex9 and TREX) share an important function in maintaining genome integrity in jawed vertebrates as “brakes” preventing LINE-1 propagation.

In studying the evolution of the PML gene, our work has implication for our understanding of how non-membrane bound organelles may have evolved in eukaryotic cells. The origins of eukaryote organelles are largely understood from the perspective of endosymbiosis. However, there are organelles that are likely derived from autogenous organellogenesis (i.e., recycling machinery and digestive vacuoles) (84). The nucleus is a complex organelle that includes a variety of subnuclear organelles within, including PML NBs, Cajal bodies, paraspeckles and splicing speckles, that contribute to its architecture and function (85,86). PML NBs are membrane-less and represent molecular condensates that self-organize into organelles, in a process regulated by SUMOylation and likely occurring by phase separation (79,87,88). PML NBs emerged in amniotes coincident with the appearance of TREX1. Amniote PML neofunctionalization appears to have occurred from the loss of exonuclease function and gain of new self-organization function primed by SUMOylation and its localization in the nucleus. Importantly, our work highlights how membrane-less organelles not derived from endosymbiosis could originate from a single gene encoding a protein with the capacity for self-organization such as PML, which has led to the amniote innovation of the PML NB.

The cellular roles of cytoplasmic PML have been relatively understudied, particularly PML-I whose role in the cytoplasm has remained a mystery for over a decade since its first observation by Condemine and colleagues (42). We discovered that L1 retroelement activation results in reproducible shuttling of PML-I to the cytoplasm in a manner that requires the NES of PML and is CRM1-dependent. Furthermore, ray-finned fish, turtle and avian PML orthologs which localize primarily and partially to the cytoplasm (respectively), can robustly inhibit L1 activity, indicating that this cytoplasmic function of PML was conserved over the course of amniote evolution. PML has also been reported to localize to the cytoplasm in response to HIV-1 infection and during epithelial-mesenchymal transition (EMT) (82,89). It will be of interest to determine if in these contexts PML nucleocytoplasmic shuttling is also CRM1-dependent and/or related to L1 activity.

The widely distributed DEDDh exonucleases play important roles in DNA replication, DNA repair, RNA maturation, and RNA turnover (90-93). Here, we describe and characterize the teleost Plex9 proteins as a novel subfamily of DEDDh exonucleases that have converged on a similar function in L1 suppression and micronuclei clearance to TREX1. We also found that while amniote PML orthologs do not encode a DEDDh exonuclease, that ray-finned Pml (sgPml) is an active exonuclease. Importantly, we demonstrate that the Plex9 proteins are not evolutionarily related to the emergence of TREX1 orthologs in amniotes, rather they are most likely derived from a separate progenitor PML-like exonuclease. Thus, the shift in PML function from exonuclease to a scaffold-like protein underlying PML NB formation was likely a consequence of redundancy in exonuclease function with the emergence of TREX1.

Our work also has implications for our understanding of the coevolution of retrotransposons and host restriction factors that restrict them in vertebrate species. While L1s contribute to genetic variation, this is juxtaposed to their negative impact on genome integrity (22,94). As a result, several mechanisms have evolved to safeguard genome stability by preventing L1 propagation, including their suppression by TREX1 (55,74). In lobe-finned vertebrates, including humans, TREX1 appears to be a major suppressor of L1s. However, prior to the emergence of TREX1, PML and Plex9 proteins have been important contributors to safeguard genomes from L1 propagation in fish. The regulation of the highly conserved cGAS-STING pathway appears to be tied to how host factors restrict or enhance L1, which has implications for innate immune signalling and senescence (25,29,74). Thus, DEDDh exonucleases have a unifying role in suppressing L1 retrotransposition that, surprisingly, is also independent of their exonuclease function.

While PML and TREX1 are both well-studied, their dynamic subcellular localization is less understood, and our study raises the question of why amniote PML retains the ability to suppress L1 elements? We found that TREX1 and PML co-localize in the cytoplasm when L1 elements are overexpressed. This leads us to conjecture that PML and TREX1 are mechanistically connected and contribute cooperatively to genome stability by suppressing retroelements, potentially via common SUMO-dependent protein interactions. Overall, our study has uncovered the convergent evolution of the Plex9 exonucleases in the teleost fish lineage that lacks TREX1 and PML orthologs, and a primordial role for PML in the suppression of L1 retrotransposition that is conserved across 350 million years of evolution.

## DATA AVAILABILITY

All genomes and transcriptomes used in the study for identifying homolog sequences are available from the National Center for Biotechnology Information (NCBI). Accession numbers and identities for sequences utilized are listed in the Materials and Methods section.

## SUPPLEMENTARY DATA

Supplementary Data is available in a separate file.

## ACKNOWLEDGEMENT

We thank Brett Racicot and Andrew W. Thompson (Michigan State University) for raising and sampling gar tissues. We also thank Dr. Nika Lovšin (University of Ljubljana) and Dr. Aurelien Doucet (CNRS) for providing zebrafish LINE-2 encoding plasmids and tagged LINE-1 ORF plasmids, respectively.

