## Supplemental Figures for "PML and PML-like exonucleases restrict retrotransposons in jawed vertebrates"

### Supplementary Figure S1

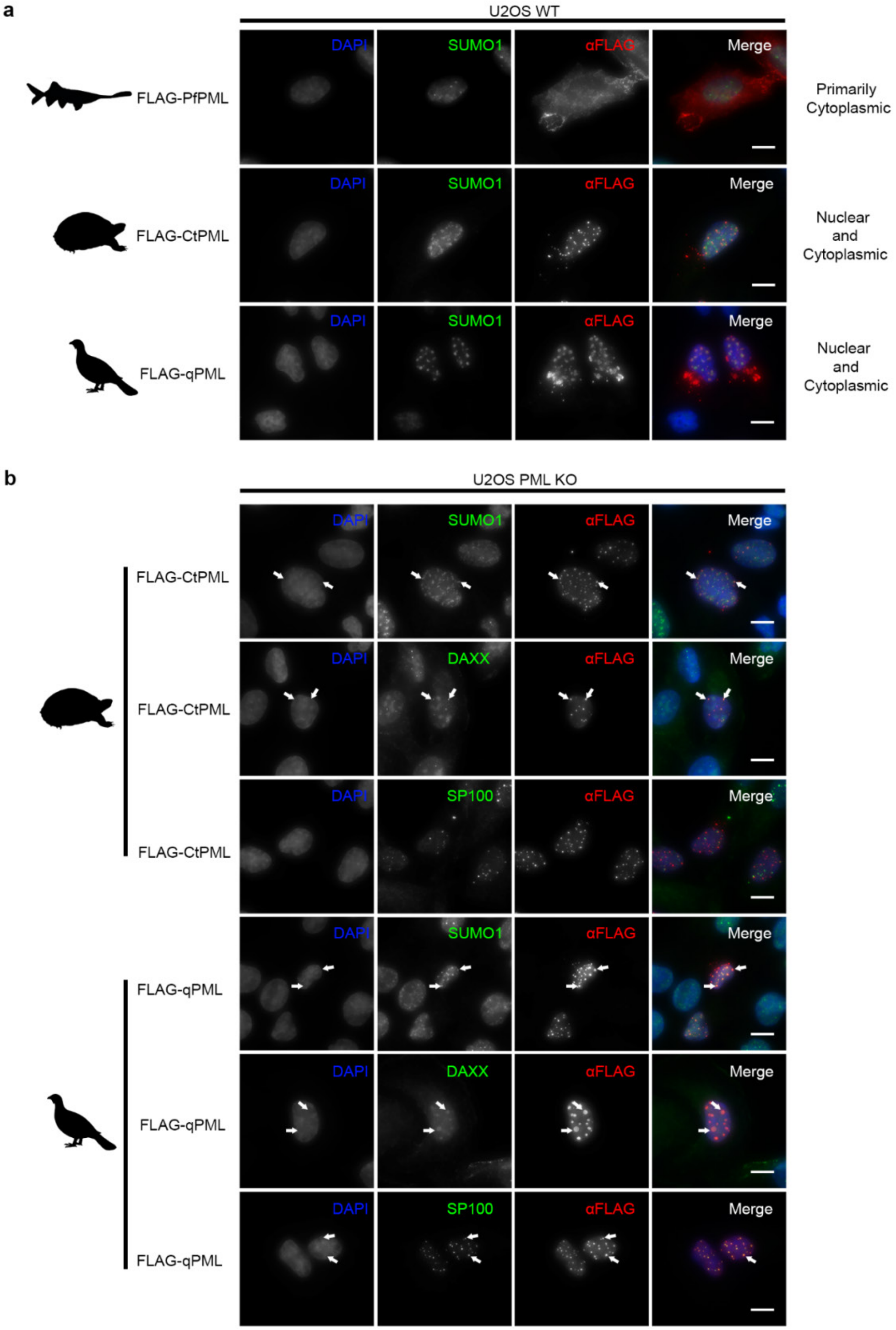


**Figure S1. Localizations of paddlefish, turtle and quail PML in human cells.** (a) Subcellular localizations of FLAG-tagged PML orthologs from American paddlefish (*Polyodon spathula*; pfPML), Common box turtle (*Terrapene Carolina*; CtPML) and Japanese quail (*Coturnix japonica;*qPML). Proteins were ectopically expressed in U2OS cells. (b) In the absence of PML (U2OS PML KO cells), turtle and quail PML form nuclear bodies that recruit DAXX and SP100, two canonical PML nuclear body components. CtPML and qPML also co-localize with SUMO. Scale bars represent 10 µm for images in (a) and (b). Arrows indicate co-localization between canonical PML NB proteins (SUMO1, DAXX and SP100) and qPML/CtPML NBs. Silhouettes for species were obtained from PhyloPic (http://phylopic.org).

### Supplementary Figure S2

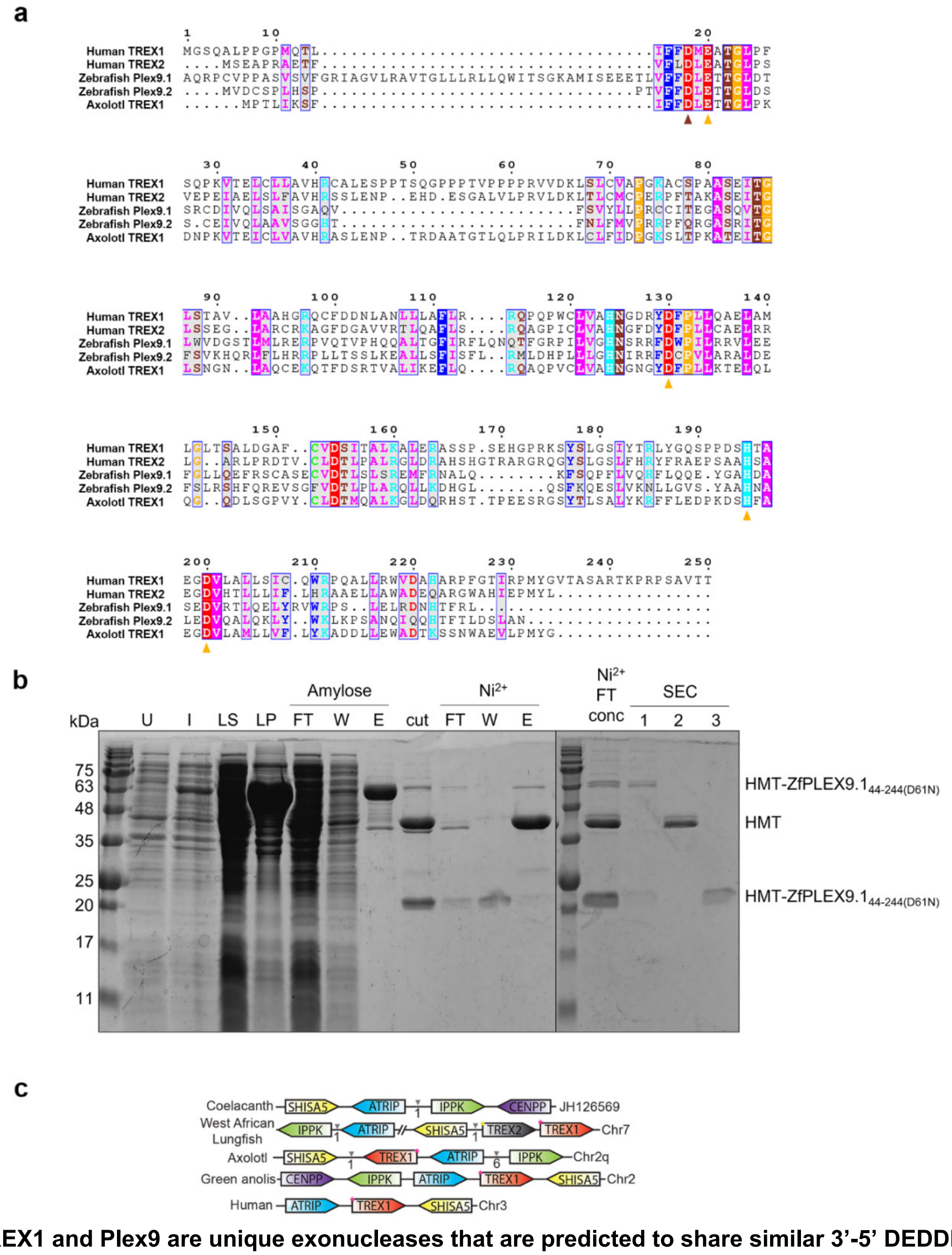


**Figure S2. TREX1 and Plex9 are unique exonucleases that are predicted to share similar 3'-5' DEDDh catalytic residues.** (a) Syntenic analysis shows that unlike Plex9, TREX1 resulted from a gene duplication of TREX2 within lobe-finned fish. (b) MUSCLE alignment of the TREX1, Plex9 and axolotl TREX1 sequences. Consensus sequences are displayed below the sequences for residues with over 70% conservation across species. Brown and yellow arrows indicate predicated catalytic residues for the 3'-5' DEDDh exonuclease domain. Brown arrow indicates the catalytic residue mutated to assess the conservation of catalytic residues**.** (c) Purification of zebrafish zfPlex9.1 (D61N). zfPlex9.1 (D61N) was expressed as a fusion to sequences encoding for hexahistidine, maltose binding protein, and a TEV protease cleavage site (HMT). SDS-PAGE analysis is shown for samples of uninduced (U) and induced (I) cells, soluble lysate (LS), after TEV cleavage (cut), flowthrough (FT), wash (W), and elution (E) fractions of amylose and $Ni^{2+}$ affinity chromatography, and fractions from size exclusion chromatography (SEC).

### Supplementary Figure S3

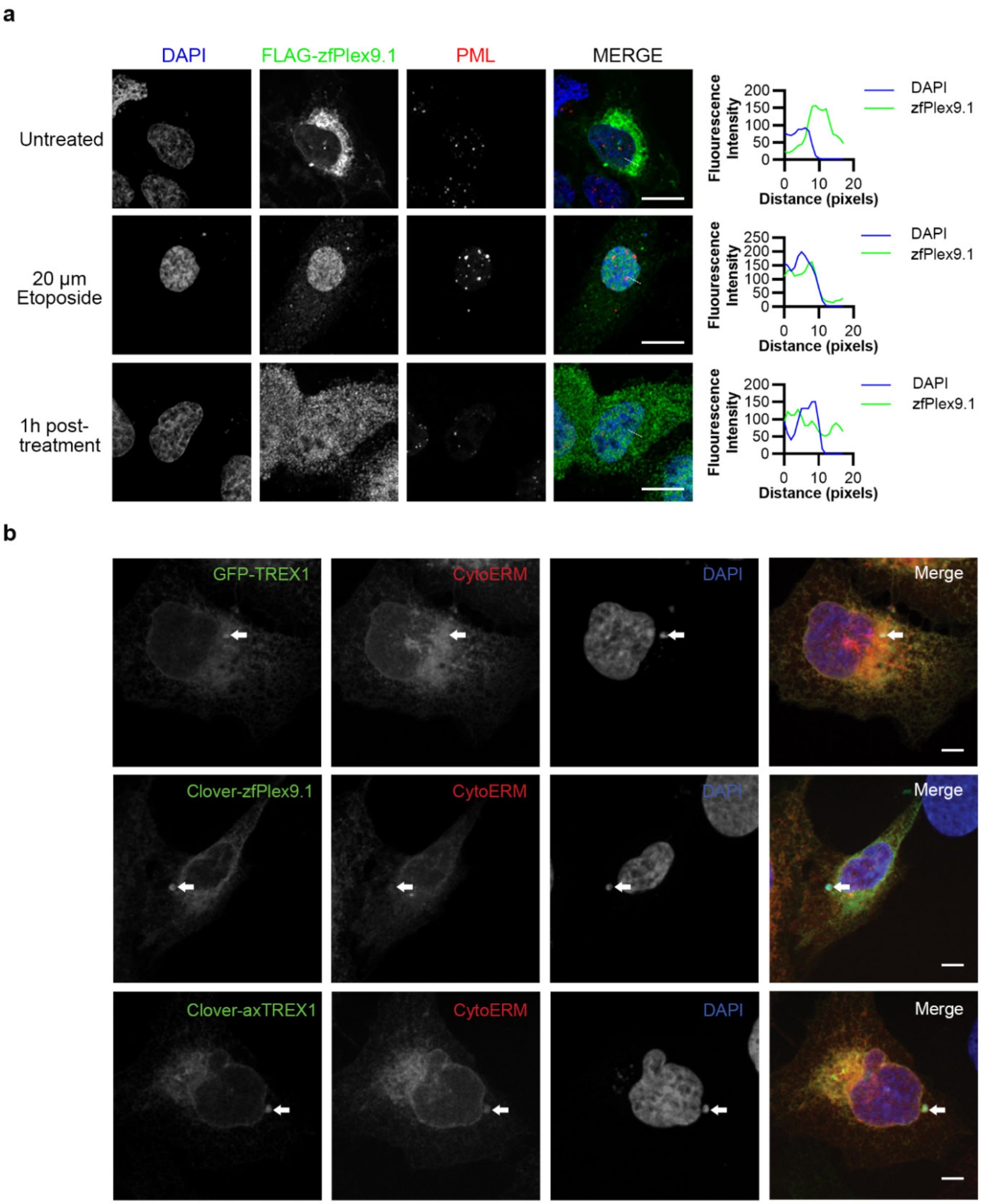


**Figure S3. Additional subcellular localizations of zfPlex9.1 and axTREX1 after DNA damage and micronuclei formation.** (a) zfPlex9.1 accumulates in the nucleus after DNA damage. FLAG-tagged zfPlex9.1 was ectopically expressed in U2OS cells. Cells were treated with etoposide (VP16 inhibitor) to induce DNA damage, 24 hours post-transfection. (b) Zebrafish zfPlex9.1 and axolotl axTREX1 accumulate at micronuclei like human TREX1. U2OS WT cells were transfected with GFP and Clover tagged human TREX1, zfPlex9.1 and axTREX1. domain of Cytochrome p450 (CytERM) tagged to mScarlet (Addgene #85066) was utilized as an ER marker to visualize ER-tubule invasion into the micronuclei. U2OS cells were treated 24 hours post-transfection with 5 µM reversine (MPS1 inhibitor used to induce micronuclei formation) and were then fixed for immunostaining Scale bars represent 5 µm for images in (a) and (b).

### Supplementary Figure S4

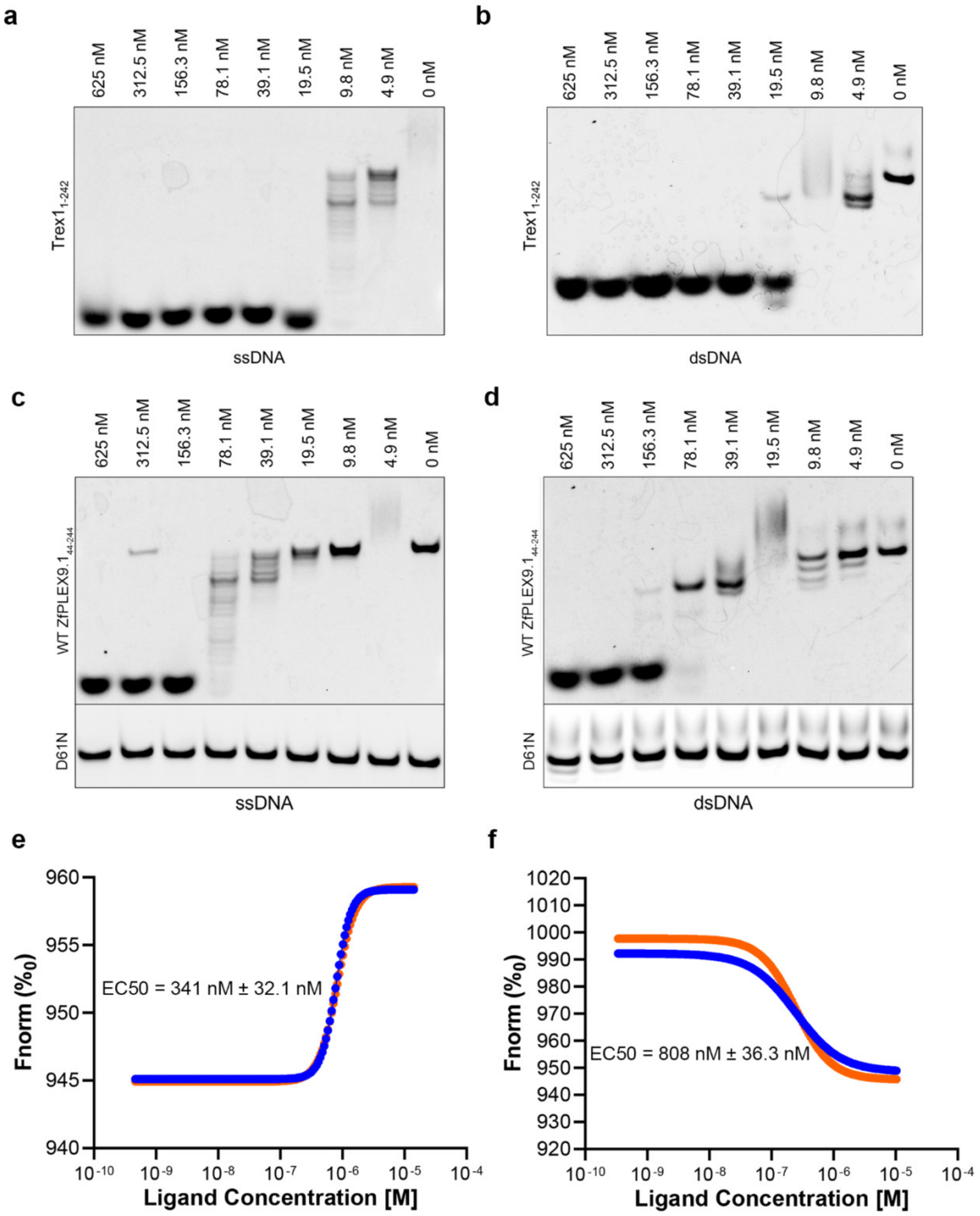


**Figure S4. The D61 residue is required for zfPlex9 enzymatic activity.** (a) and (b) Mouse Trex1 was purified akin to zfPlex9.1 and enzymatic activity was assessed for comparison of activity to zfPlex9.1. Both ssDNA (a) and dsDNA (b) were utilized as substrates at different concentrations. (c) and (d) zfPlex9 (D61N) lacks catalytic activity towards DNA. Substrates of ssDNA (c) and dsDNA (d) were utilized. Fluorescent oligonucleotides were incubated with zfPlex9.1 (D61N). Samples were resolved by UREA-PAGE and visualized via fluorescence imaging. (e) and (f) DNA binding of zfPlex9.1. Microscale thermophoresis was used to quantify the affinity between zfPlex9.1 (D61N) with (e) ssDNA ($EC_{50}$ = 341 nM ± 32.1 nM) and (f) dsDNA ($EC_{50}$ = 808 nM ± 36.3 nM) (n=2). The different coloured lines are data shown for two replicates.

### Supplementary Figure S5

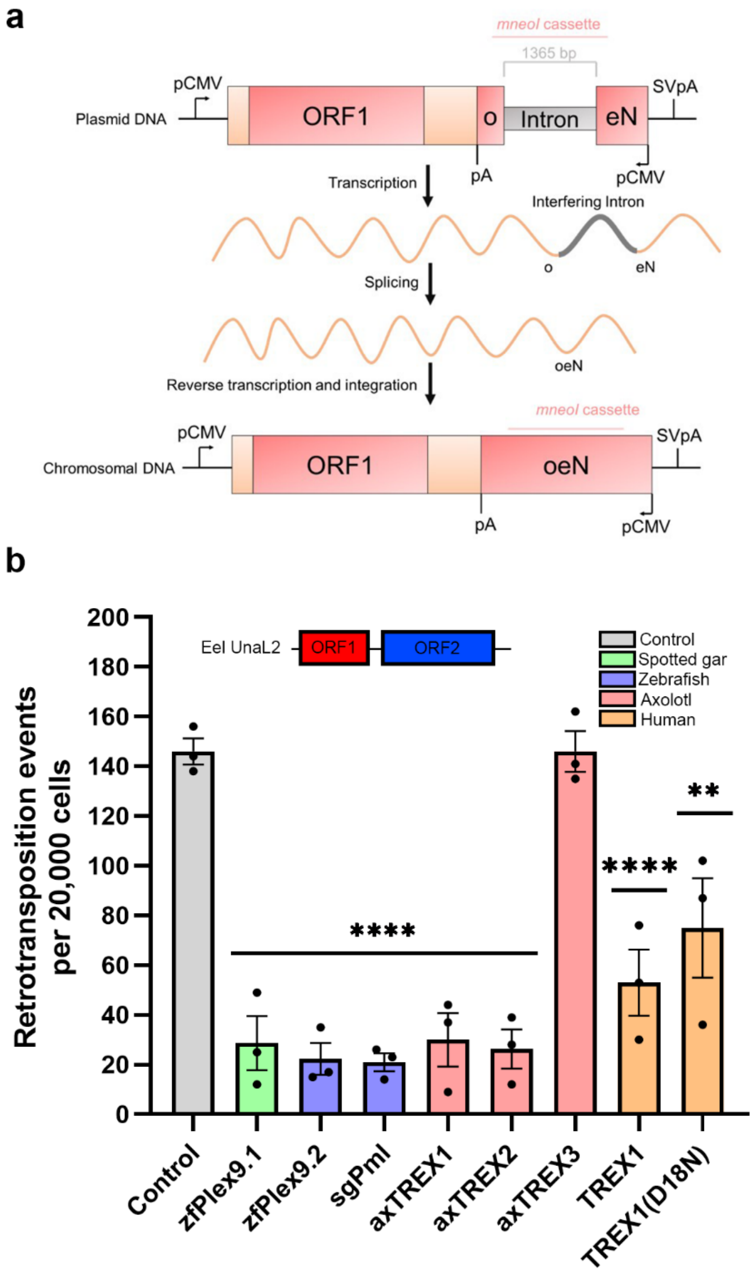


**Figure S5. UnaL2 is suppressed by Plex9, sgPml and axTREX1/2.** (a) Overview of the L1 retrotransposition assay. The LINE element adjacent to the *neo* resistance gene is present with an intervening intron. The plasmid is transfected into cells and the intervening intron is spliced out, with the *neo* gene being integrated into the chromosome of a cell with active LINEs. Co-transfection with proteins of interest allows one to study enhancers and suppressors of LINE elements. (b) Retrotransposition of human L1 was assessed with the co-expression of the FLAG-tagged versions of indicated proteins (n=3). UnaL2 is a LINE-2 isolated from eel that differs from zebrafish LINE-2 elements by its unique 3'-UTR.

### Supplementary Figure S6

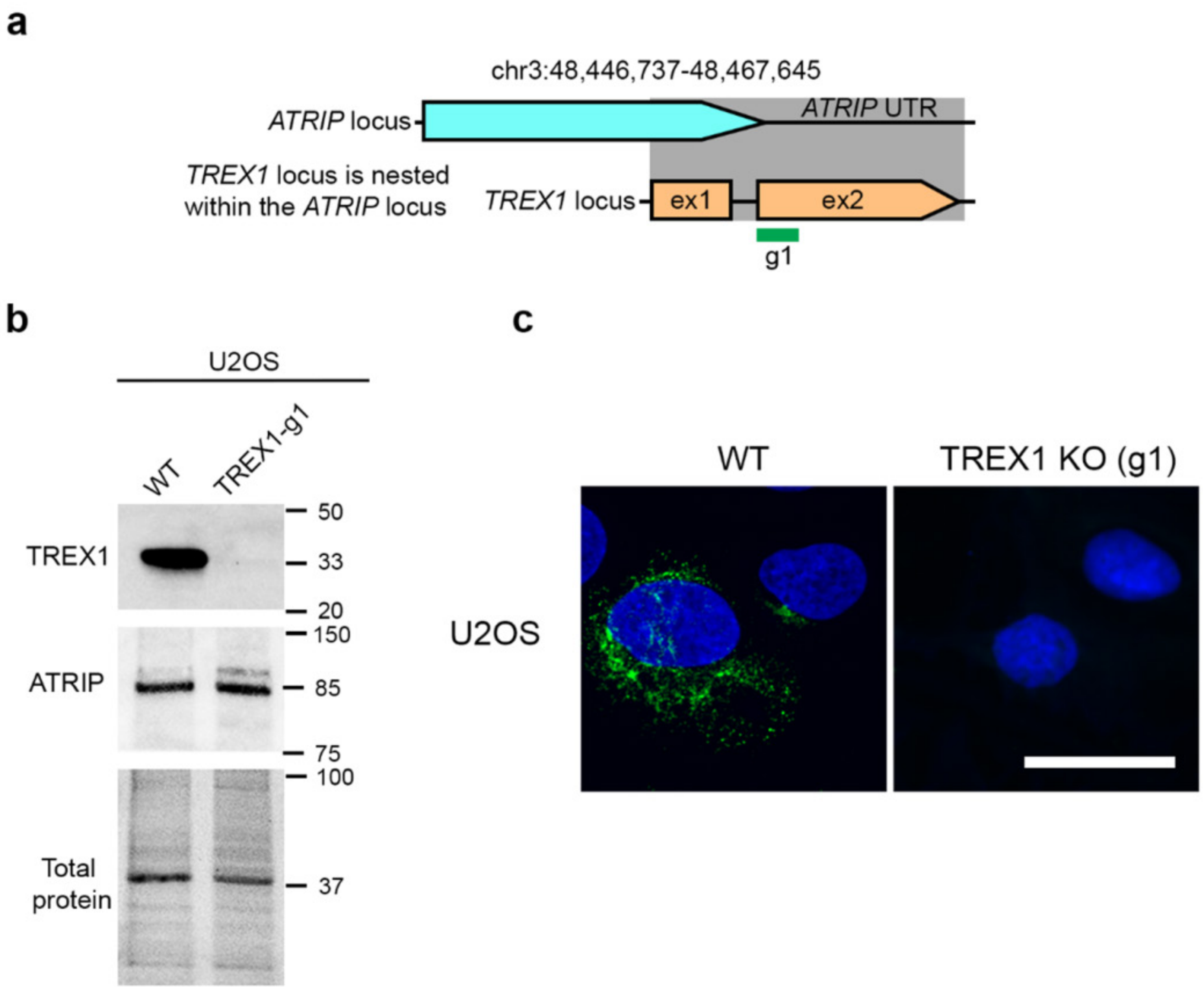


**Figure S6. Generation of a *TREX1* knockout using CRISPR/Cas9 in U2OS.** (a) Scheme for targeting human *TREX1.* A single guide RNA (sgRNA) targeting *TREX1* exon 2 (g1) was used to generate a TREX1 knockout cell line, which lacked TREX1 protein (b). Editing at the *TREX1* locus presents difficulty due to the *TREX1* gene is nested within the ATRIP UTR region and therefore we also probed for any changes in ATRIP levels. A detailed description of the CRISPR/Cas9 targeting of *TREX1* is described in the Materials and Methods. (c) Immunostaining for TREX1 in WT and *TREX1* knockout cells. Scale bar represents 20 μM.

**Supplementary Figure S7**

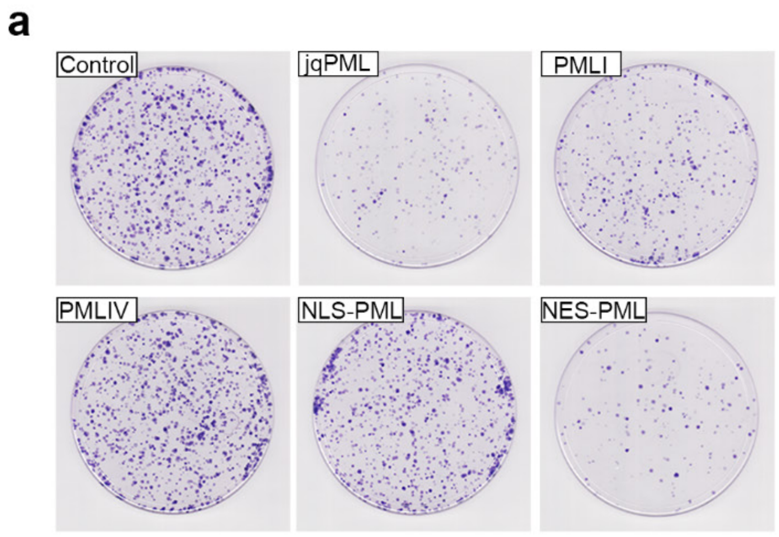


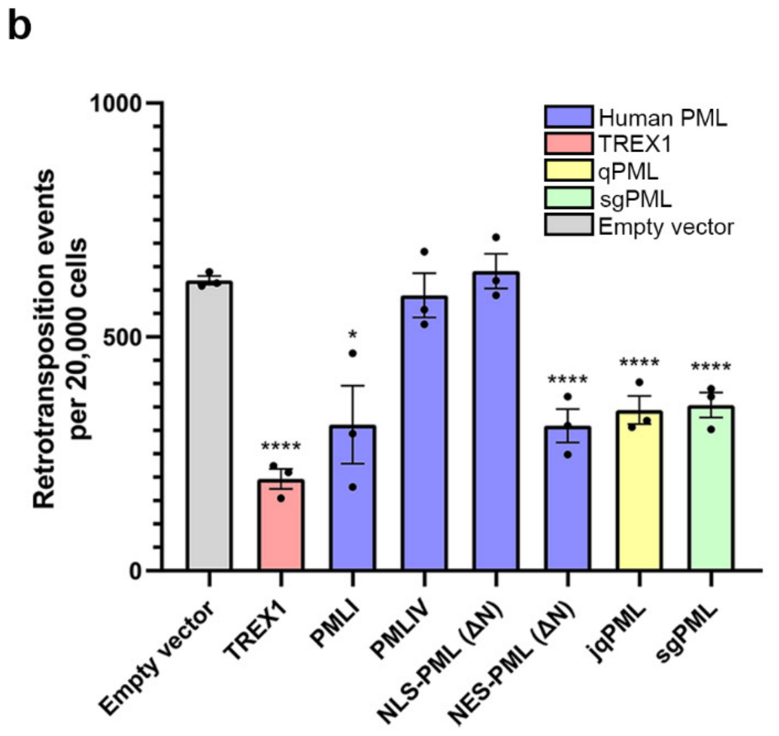


**Figure S7. PML-I and NES-PML overexpression suppresses LINE-1 in HeLa cells.** PML-I suppressed L1 activity in HeLa cells co-transfected with human L1 and the proteins described in Figure 6a. Representative plates (a) from the assay were stained with 0.5% crystal violet and quantified.

### Supplementary Figure S8

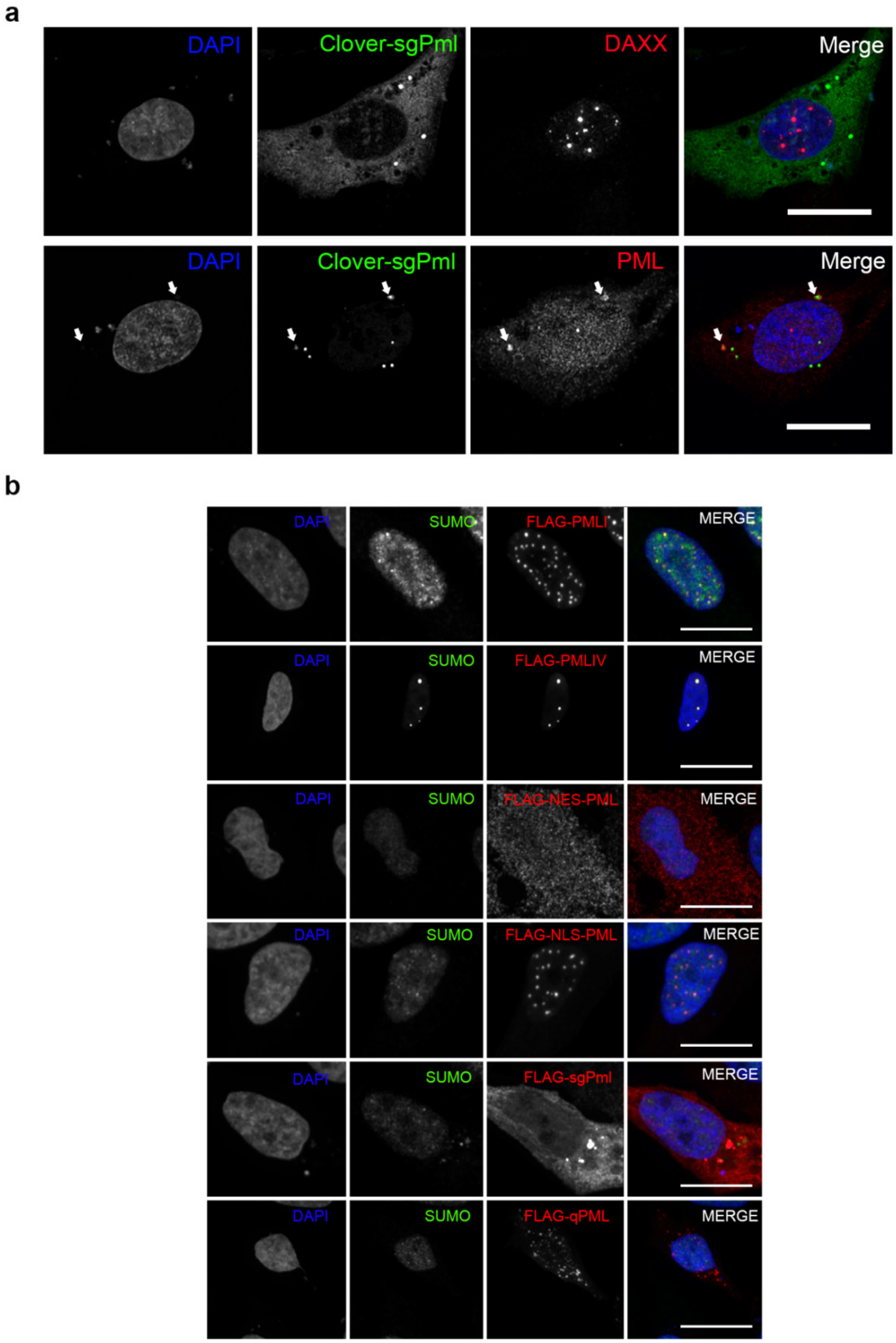


**Figure S8.** (a) Localizations of FLAG-tagged PML isoforms, mutants and orthologs described in Figure 6. (b) Co-localization of sgPml and human PML in the cytoplasm. In a small population of "stressed" cells with micronuclei, human PML and sgPml can be observed to be co-localizing in the cytoplasm. Scale bars represent 10 μM for (a) and (c).
